# Somatic mutation detection: a critical evaluation through simulations and reanalyses in oaks

**DOI:** 10.1101/2021.10.11.462798

**Authors:** Sylvain Schmitt, Thibault Leroy, Myriam Heuertz, Niklas Tysklind

## Abstract

1. Mutation, the source of genetic diversity, is the raw material of evolution; however, the mutation process remains understudied, especially in plants. Using both a simulation and reanalysis framework, we set out to explore and demonstrate the improved performance of variant callers developed for cancer research compared to single nucleotide polymorphism (SNP) callers in detecting de novo somatic mutations.
2. In an *in silico* experiment, we generated Illumina-like sequence reads spiked with simulated mutations at different allelic fractions to compare the performance of seven commonly-used variant callers to recall them. More empirically, we then reanalyzed two of the largest datasets available for plants, both developed for identifying within-individual variation in long-lived pedunculate oaks.
3. Based on the *in silico* experiment, variant callers developed for cancer research outperform SNP callers regarding plant mutation recall and precision, especially at low allele frequency. Such variants at low allelic fractions are typically expected for within-individual de novo plant mutations, which initially appear in single cells. Reanalysis of published oak data with Strelka2, the best-performing caller based on our simulations, identified up to 3.4x more candidate somatic mutations than reported in the original studies.
4. Our results advocate the use of cancer research callers to boost de novo mutation research in plants, and to reconcile empirical reports with theoretical expectations.

This version of the article has been peer-reviewed and recommended by *Peer Community in Genomics* https://doi.org/10.24072/pci.genomics.100024

## Introduction

DNA sequence mutation is the raw material for evolutionary change, but, despite its crucial role, many fundamental questions around the mutation process are still open. Notwithstanding its apparent simplicity, the understanding of mutation processes is one of the most common conceptual difficulties for biologists (Smith & Knight, 2012; Prevost et al., 2013). Mutations are often assumed to occur at a relatively constant pace (i.e. following the hypothesis of a ’perfect’ molecular clock). Despite the extremely low number of direct mutation rates estimates currently available in the literature, mutation rates are, however, known to be highly variable across the tree of life, differing by several orders of magnitude among species and kingdoms, and are considered as an evolvable trait *per se* Lynch et al., (2016). In mammals, the somatic mutation rate varies directly with life span (Cagan et al., 2022). Mutations are assumed to be random, but the rate at which different nucleotides mutate strongly depends on the genomic context, in particular the surrounding nucleotides (Martincorena & Campbell, 2015), hereafter referred to as a mutation spectrum. The mutation spectra themselves are now believed to evolve over time (Milholland et al., 2017), even at relatively short evolutionary timescales (Harris & Pritchard, 2017). The relative contribution of DNA replication and DNA repair errors to mutation rates represents another timely evolutionary question in the field (Gao et al., 2019).

Unlike most animals that transmit to the next generation only mutations present in their germ cells (*i*.*e*. sperm and eggs), plants are expected to produce heritable somatic mutations as they grow throughout their lives, departing from the so-called Weismann’s germ plasm theory (Weismann, 1893; but see also Lanfear, 2018). As a consequence, long-lived species, such as trees, are generally assumed to accumulate more heritable mutations than short-lived species per generation (Hanlon et al., 2019). To generate new knowledge on plant mutation processes (Schoen & Schultz, 2019), several studies examined within-individual variation in long-lived trees, whose individuals can live for more than a thousand years (Schöngart et al., 2017). Two studies used the pedunculate oak (*Quercus robur*), a long-lived diploid and highly heterozygous European tree species, as a plant model to identify somatic mutations. Schmid-Siegert et al., (2017) identified 17 mutations by comparing sequencing data from two branches of a 234-year-old individual. The authors therefore argued that their results are consistent with a low mutation rate in pedunculate oak. Plomion et al., (2018) identified 46 mutations using three branches of a younger century-old individual, which is an almost 10-fold higher rate after taking the tree age difference into account. Plomion et al., (2018) also recovered these new mutations on acorn embryos collected on the same branches as those used for the *de novo* mutation identification, therefore producing empirical support for departure from Weismann’s germ plasm theory in oaks. A shared limitation of both studies is that the authors used a single variant caller for *de novo* mutation detection, without investigating beforehand the robustness of the results from the selected variant caller. The absence of a simulation work to identify the best suited detection method prior to the empirical investigations therefore represents a major limit with regards to the accuracy and completeness of the previously reported *de novo* mutations.

Variant callers are designed each for a specific purpose, with choices made by the developer on sequence read filtering, and models and thresholds of sensitivity for the output of variants. Broadly, we can distinguish 1. Single nucleotide polymorphism (SNP) callers designed to detect heterozygous sites, i.e. sites with an expected allelic fractions of 0.5 in the analysed sample, and 2. Cancer callers, designed to detect sites that are mutated in a fraction of mutated cells in a sample, *i*.*e*. with expected allelic fraction ≤0.5. When used to detect mutations, SNP callers primarily detect candidate mutations per sample against the reference genome and validate mutation robustness by comparing results between sample pairs (Fig. 1). The per-sample strategy used in SNP callers carries the risk of overlooking low-frequency mutated reads in one or more samples, which risks invalidating the mutation in the other sample. To better address the specificities of detecting low frequency mutations, the development of tools to detect mutations in humans is rapidly expanding in cancer research (Kim et al., 2018; Alioto et al., 2015), where cancer callers identify mutations by comparing two samples, one mutated and one normal sample, against the reference genome (Fig. 1). Detecting mutations in cancers is conceptually similar to detecting somatic mutations in plants, i.e., the aim is to detect mutations that potentially affect only a small fraction of the sequenced tissue. However, cancer research frequently uses very high sequencing depths (100X - 1000X), while the depth available for plants is often considerably lower (e.g., 34X for Hanlon et al., 2019; 40X for Wang et al., 2018; or 70X for Schmid-Siegert et al., 2017), bar a few exceptions (240X for Orr et al., 2019; 250X for Plomion et al., 2018; or 1000X for Watson et al., 2016). Despite the advantage of cancer callers to identify mutations, the specific challenge of detecting low frequency mutations is so far poorly addressed in plants, where SNP callers have been the most frequently used method to detect somatic mutations (Schmid-Siegert et al., 2017; Watson et al., 2016; Hanlon et al., 2019; Orr et al., 2019). To demonstrate the advantages of using cancer callers in somatic mutation detection for basic and applied plant research, we evaluated the performance of the cancer callers compared to SNP callers using biological characteristics and sequencing depth typical of plant studies.

**Figure 1:**
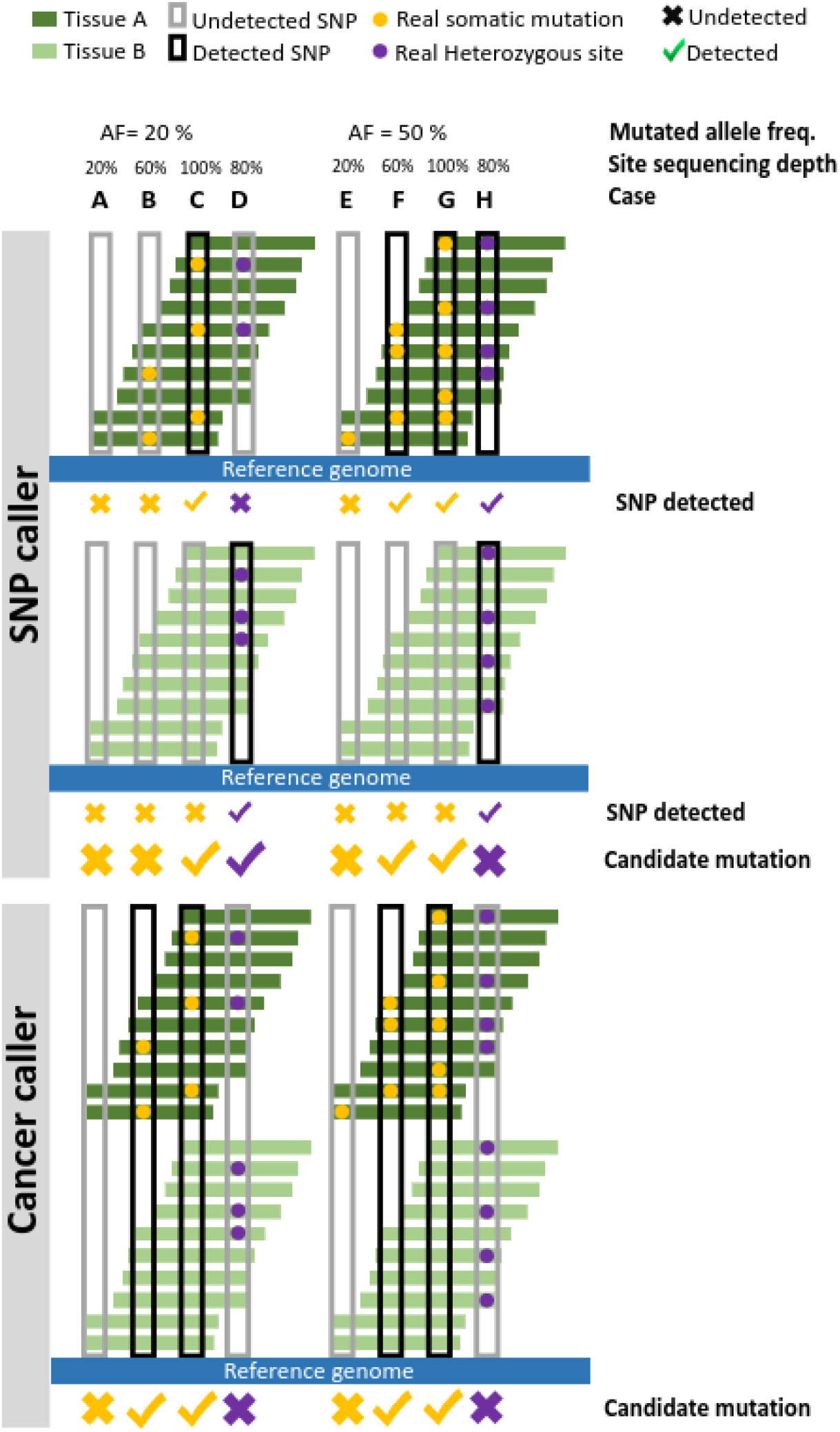
SNP callers (top rows) detect candidate mutations per tissue sample (dark green and light green) against the reference genome (blue) and validate the robustness of mutations by comparing results between sample pairs, while cancer callers identify mutations by comparing two samples, one mutated (tissue A, dark green) and one normal (tissue B, light green), against the reference genome (blue). At low sequencing depth (**A** & **E**), neither the SNP nor the cancer variant callers detect a low (**A**) or high (**E**) frequency mutation. At intermediate sequencing depths (**B** & **F**), both SNP and cancer callers detect high-frequency mutations (**F**), but cancer callers are expected to be better at detecting low-frequency mutations than SNP callers (**B**), which were originally designed to detect the expected high-frequency heterozygous sites. At high sequencing depths (**C** & **G**), both the SNP and cancer callers detect high frequency (**C**) and low frequency (**G**) mutations. However, with intermediate sequencing depth (**D** & **H**), a poorly represented heterozygous site in one tissue may remain undetected in that tissue by the SNP caller while it may be detected in the second tissue and thus be considered a mutation, resulting in a false positive (**D**). By comparing the two samples together, cancer callers will avoid this error (**H**).

Here, we performed both an *in silico* and an empirical data-based evaluation of the performance of variant callers to detect somatic mutations, using simulated reads with known mutations and two large published datasets on the same species (pedunculate oak, *Quercus robur*) that applied different strategies for sequencing depth and mutation detection (Schmid-Siegert et al., 2017; Plomion et al., 2018; see Fig. S1 (Schmitt et al., 2022)). We particularly explored the cancer callers in a plant research context to answer the following questions: (1) Can cancer research methods, both in terms of protocols (i.e. sequencing depth) and tools (i.e. callers), improve the detection of plant somatic mutations?; and (2) Can reanalyses of within-individual sequencing data provide new insights regarding plant mutation processes?

## Methods

### Study design

We developed two workflows: 1) to generate Illumina-like sequencing reads including mutations with varying biological and sequencing parameters; and 2) to detect mutations with multiple variant callers (Fig. S1). We used *singularity* containers (Kurtzer et al., 2017) and the *snakemake* workflow engines (Köster et al., 2012) to build automated, highly reproducible (FAIR), and scalable workflows. We then used both workflows to test for the best performing variant caller for mutation detection *in silico* based on biological and sequencing parameters. We finally used the identified variant caller to detect mutations in pedunculate oak, *Quercus robur* L., by re-analysing data from two somatic mutation projects on oaks led by INRAE Bordeaux, France (Plomion et al., 2018) and the University of Lausanne, Switzerland (Schmid-Siegert et al., 2017).

### Generation of mutations

To ensure the feasibility of the project and to limit the computational load, a first step is to subsample one or several sequences of user-defined length in the reference genome. The first workflow named *generateMutations* therefore uses a bespoke R script named *sample_genome* to generate these subsets (Schmitt, 2022a). The workflow then takes advantage of the two scripts included in *simuG* (v1.0.1, Yue & Liti, 2019), *vcf2model*.*pl*, and *simuG*.*pl*, respectively, 1) to build a model of heterozygous sites distribution for an haploid reference genome based on a user-defined set of known heterozygous sites in vcf format and 2) to build the second reference haploid genome comprising a user-defined number of heterozygous sites to accurately represent diploidy. Typically, the user can define a number of heterozygous sites based on the product of nucleotide diversity (π) and genome length (L). The workflow then uses a homemade R script named *generate_mutations* to spike randomly the reference genome with a user-defined number of *de novo* mutations which are drawn in a binomial distribution using a user-defined transition/transversion ratio (**R**). Finally, the workflow takes advantage of *InSilicoSeq* (v1.5.3, Gourlé et al., 2019) defined with the model option *hiseq* to generate datasets of mutated and non-mutated *in silico* Illumina-like sequencing reads with a user-defined sequencing depth (**C**), sequentially using (1) the reference haploid genome; (2) considering reference and alternate allele versions at heterozygous sites, as the workflow was developed for a diploid species; and (3) considering, or ignoring, the simulated *de novo* mutations which all feature the same user-defined allelic fraction (**AF**).

### Detection of mutations

The second workflow named detectMutations (Schmitt, 2022b) aims to detect somatic mutations from mapped sequencing reads on a genome reference. Paired-end sequencing reads of every library are quality checked using *FastQC* (v0.11.9) before trimming using *Trimmomatic* (v0.39, Bolger et al., 2014) keeping only paired-end reads without adaptors and a phred score above 15 in a sliding window of 4 bases. Reads are aligned against the reference per chromosome using *BWA mem* with the option to mark shorter splits (v0.7.17, Li & Durbin, 2009). Alignments are then compressed using *Samtools view* in CRAM format, sorted by coordinates using *Samtools sort*, and indexed using *Samtools index* (v1.10, Li et al., 2009). Duplicated reads in alignments are marked using *GATK MarkDuplicates* (v4.2.6.1, Auwera et al., 2013). Finally, the workflow uses seven variant callers to detect mutations, including variant callers developed originally for SNP calling and variant callers developed initially for cancer research. SNP callers to detect variants included *GATK HaplotypeCaller* with *GATK GenotypeGVCFs* (Auwera et al., 2013) and *freebayes* (v1.3.2, Garrison & Marth, 2012) using and reporting genotype qualities, without priors on allele balance, with a minimum alternate allelic fraction of 0.03, a minimum repeated entropy of 1 and a minimum alternate allele count of 2. Cancer callers developed for mutation detection included *VarScan* (v2.4.3, Koboldt et al., 2009), *Strelka2* (v2.9.10, Kim et al., 2018), *MuSE* (v0.1.1, Fan et al., 2016), *Mutect2* (using a panel of normal and without soft clipped bases; within v4.2.6.1; Benjamin et al., 2019), and *Somatic Sniper* (v1.0.5.0, filtering reads with mapping quality less than 25, filtering mutations with quality less than 15 with prior probability of a mutation of 0.0001; Larson et al., 2012). Then we only focused on the simulated mutations, and therefore excluded from the analyses the known heterozygous sites provided by the user thanks to the vcf file for *GATK, freebayes, Somatic Sniper*, and *Strelka2* using *BEDTools subtract* (v2.29.2, Quinlan & Hall, 2010) or directly within the variant caller for *Mutect2* and *VarScan*.

### *In silico* experiment

We used the *generateMutations* workflow to generate 1,000 mutations in the oak genome with varying biological and sequencing parameters. To ensure consistency between the *in silico* experiment and the reanalysis of empirical data, we used the reference genome “Qrob_PM1N’’ of *Quercus robur* 3P from Bordeaux, ENA accession number PRJEB8388 (Plomion et al., 2018), thus assessing the behaviour of variant callers in the same genomic context as used for the empirical work. To reduce the computational load, we only generated mutations on the first megabase of the first chromosome of the oak assembly (“Qrob_Chr01”) in order to later focus the detection on this region. To check that the conclusions regarding the callers are independent of the considered genomic region, we ran five independent investigations based on randomly selected genome areas of a megabase in length. Our results were highly congruent over all our investigations (Pearson’s correlations across all callers and all simulations, recall: 0.999, precision: 0.947, but see Table S1 for differences among callers). We used known heterozygous sites from the reference genome (Plomion et al., 2018) to simulate back ten thousand heterozygous sites (N = π x L = 10^4^, assuming π = 0.01 (Plomion et al., 2018) and L = 1 Mb). We used varying values of transition/transversion ratio (**R** = [2, 2.5, 3]), allelic fraction (**AF** = [0.05, 0.1, 0.25, 0.5]), and sequencing depth (**C** = [25, 50, 100, 150, 200]), resulting in 60 simulated datasets of mutated and associated base reads (3R x 4AF x 5C). We then used the *detectMutations* workflow to detect (recall) spiked mutations with every variant caller (*Mutect2, freebayes, GATK, Strelka2, VarScan, Somatic Sniper*, and *MuSe*). Using known spiked mutations, we assessed the number of true positive (TP), false positive (FP), and false negative (FN) for each variant caller to detect mutations and each combination of biological and sequencing parameters. We used the resulting confusion matrix to calculate the recall (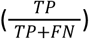) and the precision rates (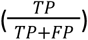). The recall rate represents the ability of the variant caller to detect all mutations, while the precision rate represents the ability of the variant caller to not confound other sites with mutations. We finally assessed each variant caller to detect mutations using the recall and the precision rates with varying transition/transversion ratio (**R**), allelic fraction (**AF**), and sequencing depth (**C**) to identify the best performing variant caller based on biological and sequencing parameters.

### Oak data reanalyses

We re-analyzed publicly available data on pedunculate oak from two projects conducted in Lausanne, Switzerland (Schmid-Siegert et al., 2017) and Bordeaux, France (Plomion et al., 2018) (SRA PRJNA327502 and ENA PRJEB8388, respectively). Sequenced reads of every library were quality checked, trimmed, and mapped with the same strategy than previously described for the simulation work, *i*.*e*. using *FastQC, Trimmomatic*, and *BWA mem*. We then used the best-performing variant caller at low coverage and allelic fraction based on our *in silico* investigation, *Strelka2*, and the variant caller for mutation detection from the original publication to compare the results, *i*.*e*., *GATK* with Best Practices for the data from Schmid-Siegert et al. (2017) and *Mutect2* for the data from Plomion et al. (2018). The former comprised 2 libraries of medium sequencing depth (60X) representing one lower and one upper branch. The latter comprised 3 libraries of high sequencing depth (160X) representing 3 branches (lower, mid, and upper). For both data sets, we compared each pair of sample points sequentially as the reference library and the potentially mutated library to distinguish mutations among branches from heterozygous sites and sequencing errors. For the data from Plomion et al. 2018, we further filtered out candidate somatic mutations by using a cross-validation procedure to keep a coherent temporal pattern among mutations following the original publication (Plomion et al., 2018). Contrary to a general expectation and a common view in the field (Schmid-Siegert et al., 2017, Orr et al., 2019), detected mutations do not always accumulate following the developing plant architecture (Zahradníková et al., 2020; Ren et al., 2021). As a consequence, our cross-validation represents a conservative strategy for the mutation detection, but it should be noted that this strategy could have removed some true somatic mutations. We used these raw datasets to identify the mutations from the original studies after realigning the megabase containing the mutation on the 3P genome using *BLAT* (Kent, 2002). For both datasets, we finally kept candidate mutations with (1) a read depth for both the normal and mutated samples between half and two times the mean sequencing depth (30-120X and 80-320X for Schmid-Siegert et al., (2017) and Plomion et al., (2018) datasets, respectively), (2) an absence of the mutated allele in the normal sample, and (3) a minimum of 10 copies of the mutated allele in the mutated sample. In addition, *Strelka2* calculates an empirical variant score (EVS) based on a supervised random forest classifier trained on data from sequencing runs under various conditions, which provides an overall quality score for each variant (Kim et al., 2018). We took advantage of the EVS to define a conservative set of candidate mutations for both datasets, hereafter referred to as the EVS datasets. Given that the proportion of the genome falling within the sequencing depth boundaries used for the detection (i.e. between 50 and 200% of the mean sequencing depth) varies depending on the dataset, we weighted the observed number of mutations by the proportion of the genome satisfying the sequencing depth criteria to provide a more accurate and comparable estimate of the real total number of mutations. Across both empirical studies, the proportion of the genome with 50-200% sequencing depth varies between 71 and 87%.

## Results

To advocate the use of cancer callers in somatic mutation detection for plant research, we simulated sequencing data containing new mutations at a given allelic fraction (*i*.*e*. fraction of simulated reads per genomic position carrying the mutated allele), and using varying depths of sequencing (for variable transition/transversion ratios, see Fig S2). We then evaluated the performance of variant callers as a function of allelic fraction and sequencing depth. We found marked differences in: (1) the recall, the ability to recover the simulated mutations; and (2) the precision, the proportion of true simulated mutations among all variants detected. For allelic fractions equal to, or lower than, 0.25, cancer variant callers (*Strelka2, Mutect2, MuSE*, but not *Somatic Sniper*) outperform SNP callers such as *GATK, freebayes*, and *VarScan* (Fig. 2 and S3), mainly based on the recall. For allelic fractions over 0.25, all variant callers perform similarly well, except for freebayes, which identified many false positives. Over the 80 tested parameter combinations, *Strelka2* was the best performing variant caller for various allelic frequencies and sequencing depths (in 57/80 simulated datasets, with an average recall of 0.95 for a precision of 0.98, Fig. S2-4 and Table S2 and S3) and an excellent computational efficiency (second fastest caller, Fig. S9).

**Figure 2:**
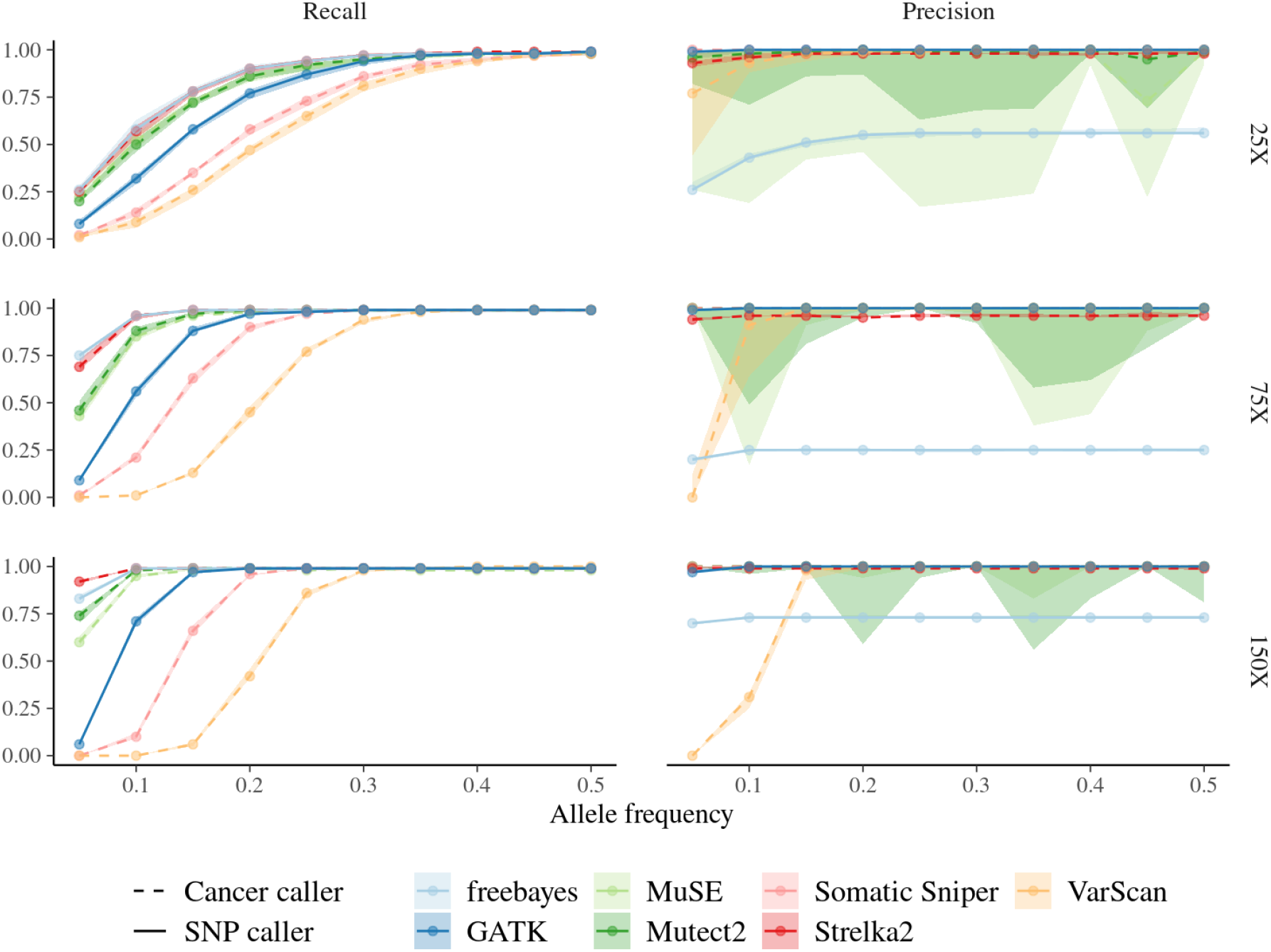
Variant caller performances to identify simulated mutations for varying allelic fractions and sequencing depths (see Fig. S5 for all parameter combinations). The recall is the ability to detect (recover) the simulated mutations. The precision is the proportion of simulated mutations among all variants detected (i.e. including false positives). Each point represents the averaged mutation recall or precision (10 simulations) for increasing allelic fraction and sequencing depth. The shaded area represents the variation of recall and precision rates over the 10 replicates computed for all callers, but only visible for the precision of *Muse, Mutect2*, and *VarScan*. Linetype distinguishes SNP callers (dashed) and cancer callers (solid).

We further investigated the performance of the best performing variant caller based on our in silico experiment, Strelka2, on two empirical datasets on pedunculate oak (Schmid-Siegert et al., 2017; Plomion et al., 2018) in comparison to the variant callers used in the original publications, i.e., GATK and Mutect2, respectively. Mapping the raw data of Schmid-Siegert et al., (2017) and Plomion et al., (2018) on the oak genome that we used as a mapping reference for our empirical study, we successfully mapped 14 and 60 of the mutations detected in the original articles, respectively. Across variant callers, we recovered 12 (86%) and 60 (100%) of these original mutations in our total list of candidate somatic mutations (Fig. 3A), strongly supporting the results shown by the two previous studies. However, our analyses were able to detect far more candidate mutations than initially reported. The filtering based on sequencing depth reduced the proportion of the genome covered with adequate sequencing depth to 72% (assuming 30-120X) for Schmid-Siegert et al., (2017) and to 84% (assuming 80-320X) for Plomion et al., (2018). Similarly, the cross-validation between branches following the plant development (see Materials and Methods for details) filtered out 27% of mutations detected for the dataset of Plomion et al., (2018). Using filtering based on sequencing depth and mutated allele copies with cross-validation, Strelka2 produced a smaller set of candidate mutations than GATK but similar to Mutect2, with an estimated number of mutation candidates 2- to 4.3-fold higher than that of the original studies (Fig. 3A). Adding Strelka2 recommended filtering based on empirical variant scores (EVS) yielded the most conservative dataset with a 1.5 to 3.4-fold increase compared to the original number of mutations. Due to lack of access to biological material from the original studies, conclusions were drawn from this list of conservative candidate somatic mutations (but see discussion regarding validation of mutations). The distribution of allelic fractions of detected mutations partly explains differences among detection methods (Fig. 3A), with Strelka2 and Mutect2 detecting mutations with lower allelic fractions than the candidate mutations presented in the original publications, especially for the Plomion et al., (2018) study that used higher sequencing depths allowing the identification of these numerous low frequency mutations.

**Figure 2:**
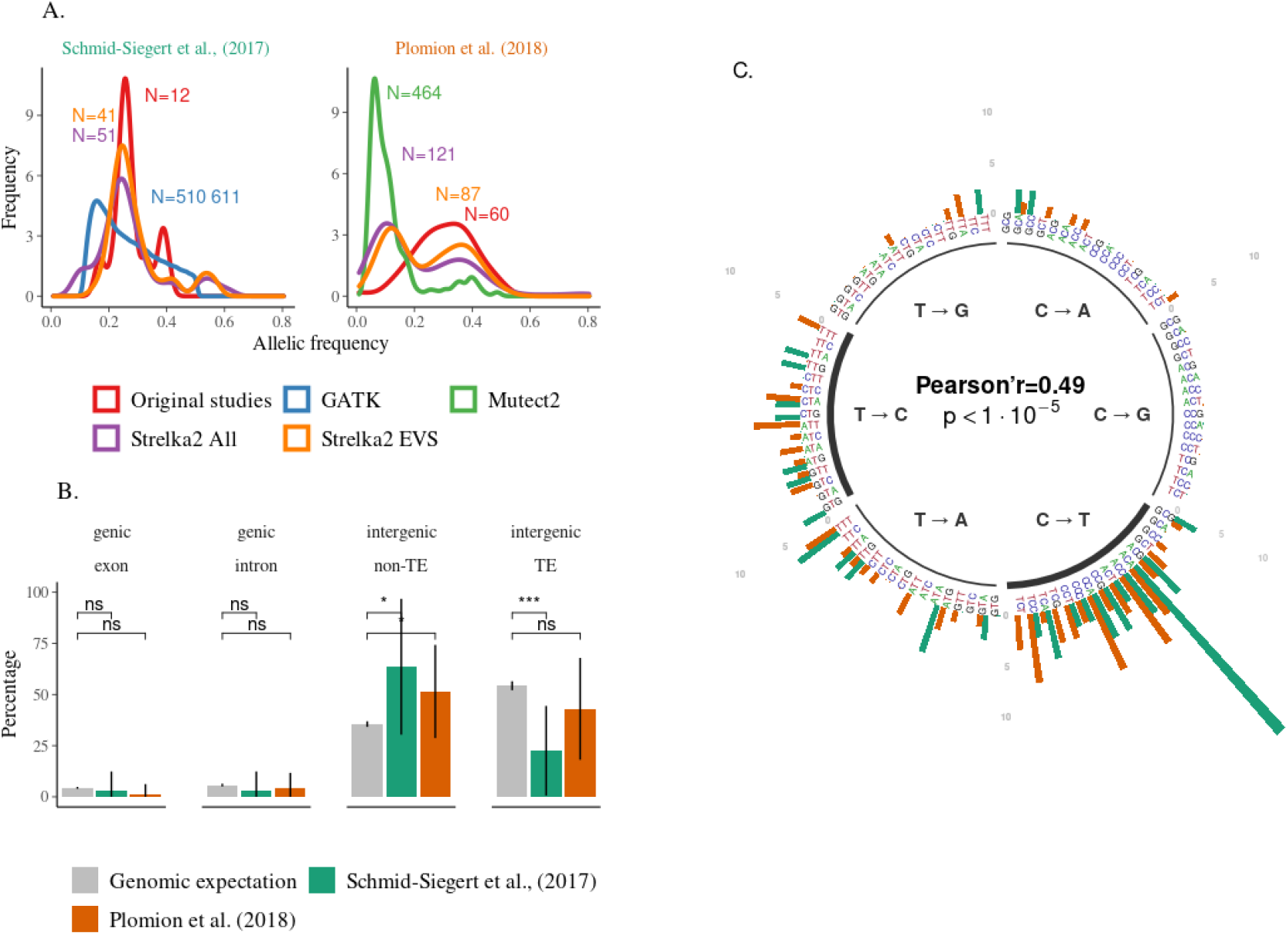
Candidate mutation spectra depending on variant callers and filtering in Schmid-Siegert et al., (2017) and Plomion et al., (2018). **A**. Allelic fraction distribution for every dataset, including the candidate mutations from the original article present in the raw data from the reanalysis (red), the results of *GATK* with Best Practices (blue), *Mutect2* after filtering (green), and *Strelka2* after filtering (purple), and the results of *Strelka2* using the filtering based on empirical variant scores named EVS (orange). The labels indicate the number of candidate mutations in each dataset. Per caller comparisons are available in Fig. S7. **B**. Annotation of the mutations detected by *Strelka2* across chromosomes using the filtering based on empirical variant scores named EVS for Schmid-Siegert et al., (2017, green) and Plomion et al., (2018, orange) compared to the genomic expectation (grey, see Supplementary Note S1). Error bars represent the standard deviation (SD) of the observed percentages across chromosomes, and the annotation above the columns indicates the significance of the Student’s t-test two-sided comparing the mean percentage of mutations to the mean genomic expectation, with ns, **, and *** corresponding to non-significant, p<0.01, and p<0.001 differences, respectively. **C**. Context-dependent mutation spectra depending on mutation types for the results of Strelka2 using the filtering based on empirical variant scores named EVS. Mutation types have been summarised into six main classes with thicker lines for transversion compared to transition, and then differentiated depending on their 5’ and 3’ genomic contexts, see Fig. S8-9. Pearson’s correlation r measures the two-sided correlation of the mutation spectra between Schmid-Siegert et al., (2017) and Plomion et al., (2018). All figures compare the reanalysed data and not the original results.

Based on the set of conservative mutations detected by *Strelka2* (EVS), we then explored annotations and mutation spectra in both datasets (Fig. 3B-C), which have rarely been explored in model plant species (but see first evidence based on mutation accumulation lines in Arabidopsis thaliana in Weng et al., 2019) and never in the wild. The proportions of mutations found in different genomic regions (e.g. genic, intergenic) were highly correlated between both original studies and proportional to the representation of the genomic regions, supporting a random distribution of mutations throughout the genome (Fig. 3B). Mutation spectra of the two studies are significantly correlated (Pearson’r=0.49, p<10-5), with an enrichment in C>T transitions, particularly in some specific genomic contexts (Fig. 3C).

## Discussion

Mutation research in plants still primarily uses SNP callers and methodologies that are not developed for the specificity and complexity of within-individual *de novo* mutation detection. We demonstrated that plant mutation research could benefit from the development of tools and protocols initially designed for human cancer research, which is a rapidly expanding field (Kim et al., 2018). We quantified expected marked differences in the performance of variant callers for mutation detection based on sequencing depth and allelic fraction. We demonstrated that cancer callers performed better than SNP callers for mutation detection at low or intermediate allelic fraction or with low sequencing depth, and similarly well for high allelic fraction. The overall higher detection efficiency demonstrates the benefit of using tools and protocols initially designed for human cancer research. Low allelic fraction mutations, potentially due to the chimeric nature of plant shoot apical meristems structures (Burian, 2021), might be very important due to their great abundance that may balance out their low chance of transmissions. Therefore, plant mutation studies should make greater use of cancer variant callers such as Strelka2 rather than SNP callers such as GATK to detect somatic mutations, in agreement with previous studies on germline mutations detection (Chen et al., 2019), especially for detecting low frequency mutations and when using low sequencing depth. The importance of allele fraction-dependency in variant detection is not restricted to somatic mutations, but also concerns for instance polyploid species, which includes many agriculturally important autopolyploid plant species (e.g. potato, sugarcane). Our simulation framework therefore provides general insights regarding the impact of allele fraction in mutation detection which go beyond somatic mutation detection.

### Validation of mutations

Intuitively, after identifying candidate mutations, one expects their “validation”. In our study, it was impossible to validate mutations using genotyping or resequencing because we performed a reanalysis of already available sequence data, without having access to the biological material. Yet, we argue that validation is more complex than generally thought. We specifically discuss (i) allele fraction, (ii) fraction-aware approach, and (iii) offspring validation. High frequency mutations (e.g. AF=0.5) will be easily confirmed by genotyping or Sanger sequencing. But if one considers that low frequency mutations are only present in a fraction of the tissue, SNP genotyping technologies are hindered by the complexity of defining the genotype clustering while Sanger sequencing is hindered by the difficulty of isolating the allelic assay from the background when reading the chromatogram. Therefore, such validation strategies are conservative and favour high frequency mutations. To unambiguously validate low-frequency mutations using intra-individual samples, an allelic fraction-aware approach can be used, such as a hybridisation capture-based sequencing strategy targeting regions of candidate mutations with high-coverage sequencing. However, this would mean that the validation is based on another round of sequencing, bringing back all the biases of the bioinformatics steps. Another strategy could be genotyping of inherited mutations in offspring, working at the family rather than the individual level, which also has its limitations (see Note S2). From our point of view, the current literature on somatic mutations in plants does not mention the limits of validation, focusing on “fully validated” mutations, thus probably favouring high frequency mutations. However, high-frequency mutations are expected to be relatively rare compared to low-frequency mutations, which affect conclusions about the total number of somatic mutations. In the relatively near future, the use of single cell sequencing approaches to reveal intra-individual genetic variation is likely to eliminate the problem at its source, avoiding the use of mixed samples of mutated and non-mutated cells, leading to simple detection and validation.

### Limits and future directions

Our study provided a first simple case of evaluating the performance of callers to detect somatic mutations in plants. Our study was limited to variant callers in the cases of mutated and normal sample pairs using the recommended default parameters. Future studies should consider and quantify the performance of other types of callers, such as subclonal or mosaic variant callers (e.g. *deepSNV* and *MosaicForecast*). Future studies could also consider exploring tuning parameters for the specific task of detecting plant somatic mutations. We hope that the open-source automated, highly reproducible, and scalable workflows constructed for our study, *generateMutations* and *detectMutations*, will allow further exploration of these questions. Our study also overlooked the importance of genome size, as plants can have larger genomes than humans. Trees have a very wide range of genome sizes, from small genomes (0.35 GB) to very large genomes (>20 GB), and our study focused on the relatively small oak genome (0.75 GB). A large genome will obviously be a barrier to high sequencing depth with a fixed budget, but with regard to mutation detection, our study argues for the use of cancer callers in large genomes as they also outperform SNP callers in terms of computational time (Fig. S9). Similarly, our study did not take into account the role of polyploidy as we only focused on a diploid species. We assume that for ploidy greater than two, the performance of cancer callers relative to SNP callers will increase since the cancer caller will be even more efficient at low frequencies. But future studies could also take advantage of our tools to explore the role of polyploidy in caller performance. Another problem that may arise when analysing sample pairs with cancer callers is the rapid increase in pairwise comparisons when using a larger sample size than previous studies (e.g., N=3 in Plomion et al. 2018). A simple solution is the use of a single reference sample such as a cambium sample from the base of the tree, which is therefore considered as the closest genome to the seed, to compare it to all samples from branches (Hanlon et al., 2019).

### Conclusion

By reanalyzing the raw oak data (Schmid-Siegert et al., 2017; Plomion et al., 2018), we found that the marked differences in the performance of variant callers could account for the discrepancies in genome-wide plant somatic mutation rate estimates. Our reanalysis using the best-performing caller based on our simulations, *Strelka2*, suggests an up to 3.4-fold higher number of mutations than previously reported, a value closer to the expectations based on the theory (Schoen & Schultz, 2019; Burian, 2021). We argue that knowledge and methodological transfers from cancer to plant mutation detection are expected to contribute strongly to the upward trend of this field and to reconcile empirical reports with theoretical expectations.

## Acknowledgements

The manuscript benefited from the comments of Nicolas Bierne and five anonymous reviewers. Preprint version 4 of this article has been peer-reviewed and recommended by Peer Community In Genomics (https://doi.org/10.24072/pci.genomics.100024).

## Data, scripts, code, and supplementary information availability

Reanalyzed reads and corresponding genomes were extracted from GenBank under accession BioProject PRJNA327502 and from European Nucleotide Archive under project accession code PRJEB8388. *generateMutations* and *detectMutations* pipelines are available online: https://doi.org/10.5281/zenodo.7274868 and https://doi.org/10.5281/zenodo.7274872. Supplementary materials are available online: https://doi.org/10.5281/zenodo.7274948.

## Authors’ contributions

All authors conceived the ideas; SS developed the pipelines, conducted the virtual experiment and the data reanalyses; SS analysed outputs and led the writing of the manuscript. All authors contributed critically to the drafts and gave final approval for publication.

## Appendix

Supplementary material is available online: https://doi.org/10.5281/zenodo.7274947.

The following Supporting Information is available for this article:

**Note S1**. Genomic expectations for the annotation of the mutations

**Note S2**. ‘Validation’ of mutations in offsprings

**Fig. S1**. Study scheme

**Fig. S2**. Variation in the performance of variant callers for mutation detection with varying biological and sequencing parameters

**Fig. S3**. Variation in the performance of variant callers for mutation detection with varying biological and sequencing parameters

**Fig. S4**. Best performing variant callers for mutation detection depending on allelic fraction (allelic fraction) and coverage (sequencing depth)

**Fig. S5**. Mutation recall and precision rates for SNP and cancer variant callers by allelic fraction and sequencing depth

**Fig. S6**. Observed allelic frequencies of candidate mutations depending on variant callers and filtering in Schmid-Siegert et al., (2017) and Plomion et al., (2018)

**Fig. S7**. Percentage of nucleotide change types of candidate mutations depending on variant callers and filtering in Schmid-Siegert et al., (2017) and Plomion et al., (2018)

**Fig. S8**. Context-dependent mutation spectrum depending on variant callers and filtering in Schmid-Siegert et al., (2017) and Plomion et al., (2018)

**Tab. S1**. Mean and standard deviation in performance of variant callers for mutation detection across all simulations

**Tab S2**. Mean and standard deviation in performance of variant callers for mutation detection with varying allelic fraction and sequencing depth

## References

Alioto, TS, Buchhalter I, Derdak S, Hutter B, Eldridge MD, Hovig E, Heisler LE, Beck TA, Simpson JT, Tonon L, Sertier AS, Patch AM, Jäger N, Ginsbach P, Drews R, Paramasivam N, Kabbe R, Chotewutmontri S, Diessl N, Previti C, Schmidt S, Brors B, Feuerbach L, Heinold M, Gröbner S, Korshunov A, Tarpey PS, Butler AP, Hinton J, Jones D, Menzies A, Raine K, Shepherd R, Stebbings L, Teague JW, Ribeca P, Giner FC, Beltran S, Raineri E, Dabad M, Heath SC, Gut M, Denroche RE, Harding NJ, Yamaguchi TN, Fujimoto A, Nakagawa H, Quesada V, Valdés-Mas R, Nakken S, Vodák D, Bower L, Lynch AG, Anderson CL, Waddell N, Pearson JV, Grimmond SM, Peto M, Spellman P, He M, Kandoth C, Lee S, Zhang J, Létourneau L, Ma S, Seth S, Torrents D, Xi L, Wheeler DA, López-Otín C, Campo E, Campbell PJ, Boutros PC, Puente XS, Gerhard DS, Pfister SM, McPherson JD, Hudson TJ, Schlesner M, Lichter P, Eils R, Jones DTW, Gut IG (2015). A comprehensive assessment of somatic mutation detection in cancer using whole-genome sequencing. Nature Communications, 6. https://doi.org/10.1038/ncomms10001

Auwera GA, Carneiro MO, Hartl C, Poplin R, Angel G, Levy-Moonshine A, Jordan T, Shakir K, Roazen D, Thibault J, Banks E, Garimella KV, Altshuler D, Gabriel S, DePristo MA (2018) From FastQ Data to High-Confidence Variant Calls: The Genome Analysis Toolkit Best Practices Pipeline. Current Protocols in Bioinformatics, 43, 483–492. https://doi.org/10.1002/0471250953.bi1110s43

Benjamin D, Sato T, Cibulskis K, Getz G, Stewart C, Lichtenstein L (2019) Calling Somatic SNVs and Indels with Mutect2. bioRxiv. https://doi.org/10.1101/861054

Bolger AM, Lohse M, Usadel B (2014) Trimmomatic: A flexible trimmer for Illumina sequence data. Bioinformatics, 30, 2114–2120. https://doi.org/10.1093/bioinformatics/btu170

Burian A (2021) Does Shoot Apical Meristem Function as the Germline in Safeguarding Against Excess of Mutations? Frontiers in Plant Science, 12, 1–9. https://doi.org/10.3389/fpls.2021.707740

Cagan A, Baez-Ortega A, Brzozowska N, Abascal F, Coorens THH, Sanders MA, … Martincorena I (2022) Somatic mutation rates scale with lifespan across mammals. Nature, 604(7906), 517–524. https://doi.org/10.1038/s41586-022-04618-z

Chen ZL, Meng JM, Cao Y, Yin JL, Fang RQ, Fan SB, Liu C, Zeng WF, Ding YH, Tan D, Wu L, Zhou WJ, Chi H, Sun RX, Dong MQ, He SM (2019) A high-speed search engine pLink 2 with systematic evaluation for proteome-scale identification of cross-linked peptides. Nature Communications, 10. http://dx.doi.org/10.1038/s41467-019-11337-z

Fan Y, Xi L, Hughes DST, Zhang J, Zhang J, Futreal PA, Wheeler DA, Wang W (2016) MuSE: accounting for tumor heterogeneity using a sample-specific error model improves sensitivity and specificity in mutation calling from sequencing data. Genome biology, 17, 178. http://dx.doi.org/10.1186/s13059-016-1029-6

Gao Z, Moorjani P, Sasani TA, Pedersen BS, Quinlan AR, Jorde LB, Amster G, Przeworski M (2019) Overlooked roles of DNA damage and maternal age in generating human germline mutations. Proceedings of the National Academy of Sciences of the United States of America, 116, 9491–9500. https://doi.org/10.1073/pnas.1901259116

Garrison E, Marth G (2012) Haplotype-based variant detection from short-read sequencing. arXiv, 1–9. https://doi.org/10.48550/arXiv.1207.3907

Gourlé H, Karlsson-Lindsjö O, Hayer J, Bongcam-Rudloff E (2019) Simulating Illumina metagenomic data with InSilicoSeq. Bioinformatics, 35, 521–522. https://doi.org/10.1093/bioinformatics/bty630

Hanlon VCT, Otto SP, Aitken SN (2019) Somatic mutations substantially increase the per-generation mutation rate in the conifer Picea sitchensis. Evolution Letters, 3, 348–358. https://doi.org/10.1002/evl3.121

Harris K, Pritchard JK (2017) Rapid evolution of the human mutation spectrum. eLife, 6, 1–17. https://doi.org/10.7554/eLife.24284

Kent WJ (2002) BLAT—The BLAST-Like Alignment Tool. Genome Research, 12, 656–664. https://doi.org/10.1101/gr.229202

Kim S, Scheffler K, Halpern AL, Bekritsky MA, Noh E, Källberg M, Chen X, Kim Y, Beyter D, Krusche P, Saunders CT (2018) Strelka2: fast and accurate calling of germline and somatic variants. Nature Methods, 15, 591–594. https://doi.org/10.1038/s41592-018-0051-x

Koboldt DC, Chen K, Wylie T, Larson DE, McLellan MD, Mardis ER, Weinstock GM, Wilson RK, Ding L (2009) VarScan: Variant detection in massively parallel sequencing of individual and pooled samples. Bioinformatics, 25, 2283–2285. https://doi.org/10.1093/bioinformatics/btp373

Köster J, Rahmann S (2012) Snakemake-a scalable bioinformatics workflow engine. Bioinformatics, 28, 2520–2522. https://doi.org/10.1093/bioinformatics/bts480

Kurtzer GM, Sochat V, Bauer MW (2017) Singularity: Scientific containers for mobility of compute. PLoS ONE, 12, 1–20. https://doi.org/10.1371/journal.pone.0177459

Lanfear R (2018) Do plants have a segregated germline? PLoS Biology, 16, 1–13. https://doi.org/10.1371/journal.pbio.2005439

Larson DE, Harris CC, Chen K, Koboldt DC, Abbott TE, Dooling DJ, Ley TJ, Mardis ER, Wilson RK, Ding L (2012) Somaticsniper: Identification of somatic point mutations in whole genome sequencing data. Bioinformatics, 28, 311–317. https://doi.org/10.1093/bioinformatics/btr665

Li H, Durbin R (2009) Fast and accurate short read alignment with Burrows-Wheeler transform. Bioinformatics, 25, 1754–1760. https://doi.org/10.1093/bioinformatics/btp324

Li H, Handsaker B, Wysoker A, Fennell T, Ruan J, Homer N, Marth G, Abecasis G, Durbin R (2009) The Sequence Alignment/Map format and SAMtools. Bioinformatics, 25, 2078–2079. https://doi.org/10.1093/bioinformatics/btp352

Lynch M, Ackerman MS, Gout JF, Long H, Sung W, Thomas WK, Foster PL (2016) Genetic drift, selection and the evolution of the mutation rate. Nature Reviews Genetics, 17, 704–714. http://dx.doi.org/10.1038/nrg.2016.104

Martincorena I, Campbell PJ (2015). Somatic mutation in cancer and normal cells. Science, 349, 1483–1489. https://doi.org/10.1126/science.aab4082

Milholland B, Dong X, Zhang L, Hao X, Suh Y, Vijg J (2017) Differences between germline and somatic mutation rates in humans and mice. Nature Communications, 8, 1–8. http://dx.doi.org/10.1038/ncomms15183

Orr AJ, Padovan A, Kainer D, Külheim C, Bromham L, Bustos-Segura C, Foley W, Haff T, Hsieh JF, Morales-Suarez A, Cartwright RA, Lanfear R (2020) A phylogenomic approach reveals a low somatic mutation rate in a long-lived plant. Proc. R. Soc. B, 287, 20192364–20192364. http://doi.org/10.1098/rspb.2019.2364

Plomion C, Aury JM, Amselem J, Leroy T, Murat F, Duplessis S, Faye S, Francillonne N, Labadie K, Le Provost G, Lesur I, Bartholomé J, Faivre-Rampant P, Kohler A, Leplé JC, Chantret N, Chen J, Diévart A, Alaeitabar T, Barbe V, Belser C, Bergès H, Bodénès C, Bogeat-Triboulot MB, Bouffaud ML, Brachi B, Chancerel E, Cohen D, Couloux A, Da Silva C, Dossat C, Ehrenmann F, Gaspin C, Grima-Pettenati J, Guichoux E, Hecker A, Herrmann S, Hugueney P, Hummel I, Klopp C, Lalanne C, Lascoux M, Lasserre E, Lemainque, A, Desprez-Loustau ML, Luyten I, Madoui MA, Mangenot S, Marchal C, Maumus F, Mercier J, Michotey C, Panaud O, Picault N, Rouhier N, Rué O, Rustenholz C, Salin F, Soler M, Tarkka M, Velt A, Zanne AE, Martin F, Wincker P, Quesneville H, Kremer A, Salse J (2018) Oak genome reveals facets of long lifespan. Nature Plants, 4, 440–452. http://dx.doi.org/10.1038/s41477-018-0172-3

Prevost L, Knight J, Smith M, Lurain UM (2013). Student writing reveals their heterogeneous thinking about the origin of genetic variation in populations. National Association on Research in Science Teaching. https://www.colorado.edu/sei/content/student-writing-reveals-their-heterogeneous-thinking

Quinlan AR, Hall IM (2010) BEDTools: A flexible suite of utilities for comparing genomic features. Bioinformatics, 26, 841–842. https://doi.org/10.1093/bioinformatics/btq033

Ren Y, He Z, Liu P, Traw B, Sun S, Tian D, Yang S, Jia Y, Wang L (2021) Somatic Mutation Analysis in Salix suchowensis Reveals Early-Segregated Cell Lineages. Molecular Biology and Evolution, 38, 5292–5308. https://doi.org/10.1093/molbev/msab286

Schmid-Siegert E, Sarkar N, Iseli C, Calderon S, Gouhier-Darimont C, Chrast J, Cattaneo P, Schütz F, Farinelli L, Pagni M, Schneider M, Voumard J, Jaboyedoff M, Fankhauser C, Hardtke CS, Keller L, Pannell JR, Reymond A, Robinson-Rechavi M, Xenarios I, Reymond P (2017) Low number of fixed somatic mutations in a long-lived oak tree. Nature Plants, 3, 926–929. http://dx.doi.org/10.1038/s41477-017-0066-9

Schmitt S (2022). generateMutations: singularity & snakemake workflow to generate in silico mutations. Zenodo, https://doi.org/10.5281/zenodo.7274868

Schmitt S (2022). detectMutations: singularity & snakemake workflow to detect mutations with several callers. Zenodo, https://doi.org/10.5281/zenodo.7274872

Schmitt S, Leroy T, Heuertz M, Tysklind T (2022). Supplementary material of Somatic mutation detection: a critical evaluation through simulations and reanalyses in oaks. Zenodo, https://doi.org/10.5281/zenodo.7274948

Schoen DJ, Schultz ST (2019) Somatic Mutation and Evolution in Plants. Annual Review of Ecology, Evolution, and Systematics, 50, 49–73. https://doi.org/10.1146/annurev-ecolsys-110218-024955

Schöngart J, Bräuning A, Barbosa ACMC, Lisi CS, Oliveira JM (2017) Dendroecology. Tree-Ring Analyses Applied to Ecological Studies. Springer. https://doi.org/10.1007/978-3-319-61669-8

Smith MK, Knight JK (2012) Using the Genetics Concept Assessment to document persistent conceptual difficulties in undergraduate genetics courses. Genetics, 191(1), 21–32. https://doi.org/10.1534/genetics.111.137810

Wang L, Ji Y, Hu Y, Hu H, Jia X, Jiang M, Zhang X, Zhao L, Zhang Y, Jia Y, Qin C, Yu L, Huang J, Yang S, Hurst LD, Tian D (2019) The architecture of intra-organism mutation rate variation in plants. PLoS Biology, 17, 1–29. https://doi.org/10.1371/journal.pbio.3000191

Watson JM, Platzer A, Kazda A, Akimcheva S, Valuchova S, Nizhynska V, Nordborg M, Riha K (2016) Germline replications and somatic mutation accumulation are independent of vegetative life span in Arabidopsis. Proceedings of the National Academy of Sciences of the United States of America, 113, 12226–12231. https://doi.org/10.1073/pnas.1609686113

Weismann A (1893) The germ-plasm: a theory of heredity. Scribner’s. http://www.esp.org/books/weismann/germ-plasm/facsimile/

Weng ML, Becker C, Hildebrandt J, Neumann M, Rutter MT, Shaw RG, Weigel D, Fenster CB (2019) Fine-grained analysis of spontaneous mutation spectrum and frequency in arabidopsis thaliana. Genetics, 211, 703–714. https://doi.org/10.1534/genetics.118.301721

Yue, J.X. & Liti, G. (2019). SimuG: A general-purpose genome simulator. Bioinformatics, 35, 4442–4444.

Zahradníková E, Ficek A, Brejová B, Vinař T, Mičieta K (2020) Mosaicism in old trees and its patterns. Trees - Structure and Function, 34, 357–370. https://doi.org/10.1007/s00468-019-01921-7

