## Supplementary material for "Somatic mutation detection: a critical evaluation through simulations and reanalyses in oaks": Fig. S1 (from another version of this preprint: version 3, 2022-08-16)

Article title: Somatic mutation detection in plants: a critical evaluation through simulations and reanalyses

All following supplementary materials can be accessed as appendix to the main manuscript at: https://doi.org/10.1101/590836

**Note S1.** Genomic expectations for the annotation of the mutations

**Note S2.** ‘Validation’ of mutations in offsprings

**Fig. S1.** Study scheme

**Fig. S2.** Variation in the performance of variant callers for mutation detection with varying biological and sequencing parameters

**Fig. S3.** Variation in the performance of variant callers for mutation detection with varying biological and sequencing parameters

**Fig. S4.** Best performing variant callers for mutation detection depending on allelic fraction (allelic fraction) and coverage (sequencing depth)

**Fig. S5.** Mutation recall and precision rates for SNP and cancer variant callers by allelic fraction and sequencing depth

**Fig. S6.** Observed allelic frequencies of candidate mutations depending on variant callers and filtering in Schmid-Siegert *et al.,* (2017) and Plomion *et al.,* (2018)

**Fig. S7.** Percentage of nucleotide change types of candidate mutations depending on variant callers and filtering in Schmid-Siegert *et al.,* (2017) and Plomion *et al.,* (2018)

**Fig. S8.** Context-dependent mutation spectrum depending on variant callers and filtering in Schmid-Siegert *et al.,* (2017) and Plomion *et al.,* (2018)

**Tab. S1.** Correlations between simulations on fixed or random genomic areas.

**Tab. S2.** Mean and standard deviation in performance of variant callers for mutation detection across all simulations

**Tab S3.** Mean and standard deviation in performance of variant callers for mutation detection with varying allelic fraction and sequencing depth

**Note S1: Genomic expectations for the annotation of the mutations**

Gene and tandem repeat (TE) element annotations from the 3P oak genome (Plomion *et al.,* 2018) were used to calculate the genomic expectations for mutations in genomic regions according to their annotation, assuming mutations were randomly distributed on each chromosome. We identified the categories of “genic exons”, grouping the coding DNA sequence (CDS) and 5' and 3' untranslated regions (UTRs), and that of “genic introns”, that corresponded to the rest of the gene annotations. “Intergenic regions” comprised all genomic regions not annotated as genes, which included intergenic TEs also annotated as TEs and the rest being the intergenic non-TEs.

**Note S2: ‘Validation’ of mutations in offsprings**

Another strategy could be the genotyping of mutations that are passed to the progeny, so as to work at the family level for the validation rather than within-individuals. A mutation, when present in the gamete, is then present in all cells of the new individual (i.e. passed at f=0.5, assuming outcrosses), allowing the validation of the mutation by both genotyping and Sanger sequencing. Note that this approach allows the validation of the new mutations but it does not mean that it is simple, at least with a genotyping strategy, because the family of offspring is expected to contain only two genotype categories, homozygous for wild type, 0/0, and heterozygous, 0/1. However, the identification of clusters in genotyping is typically based on three categories, including 1/1. In the absence of double-mutated genotypes (i.e. no 1/1 genotypes, except assuming the selfing of two gametes of the same branch, when both carry the mutation), a lower quality of the genotyping is likely to result and, in the worst case, the removing of some genotyped loci during the analysis (Leroy, personal observation). In addition, it should be noted that this strategy allows the validation of mutations that are passed to the next generation, but other detected somatic mutations (about 50% in Plomion *et al.,* 2018 and Wang et al. 2019) could be real mutations as well, albeit not identified as transmitted using their experimental design, as interpreted by Plomion and collaborators in their paper (Plomion *et al.,* 2018). Reasons for the non-detection of real mutations in offspring could lie in the AF of the mutation in the tissue and the position of the mutated cell line (cell layer) in the apical shoot meristem, in association with the number of available/genotyped offspring.

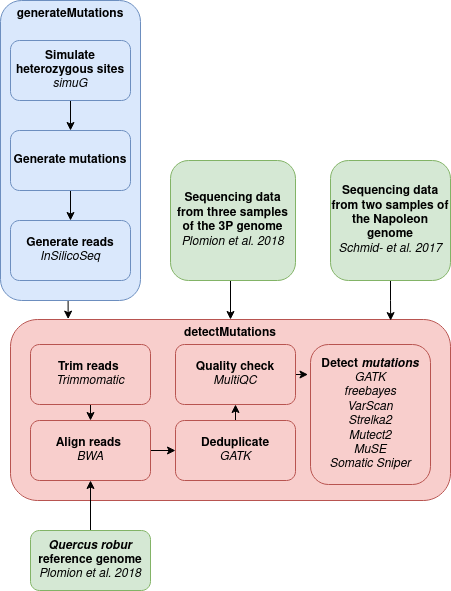

*Figure S1: Study scheme. The two highly reproducible workflows to generate Illumina-like sequencing reads and detect known mutations in simulated and publicly available empirical data sets are coloured in blue and red, respectively. Green indicated publicly available empirical data sets.*

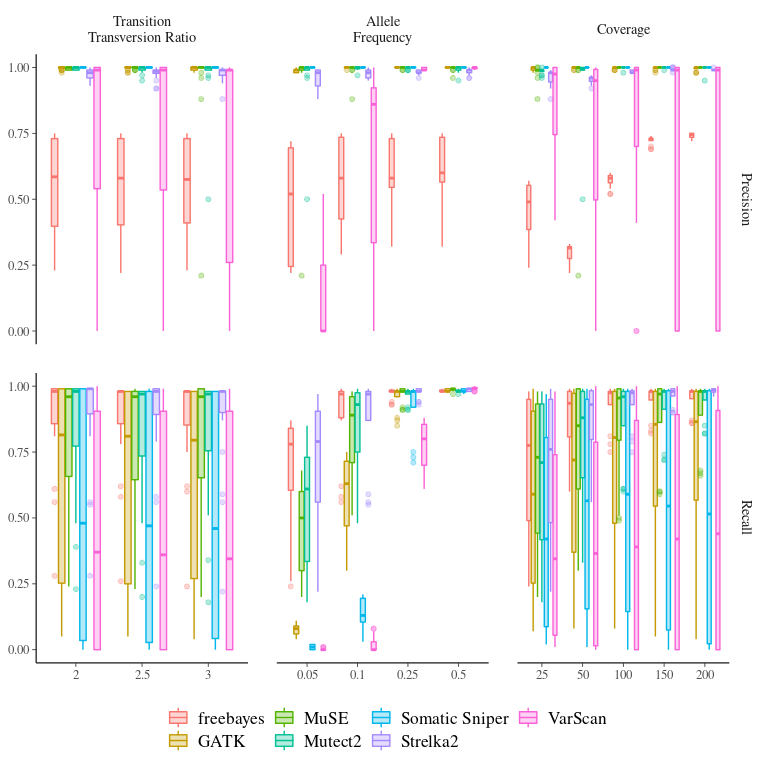

*Figure S2: Variation in the performance of variant callers for mutation detection with varying biological and sequencing parameters. The recall and the precision rates have been assessed for each variant caller to detect mutations with varying transition/transversion ratio, allelic fraction, and coverage (sequencing depth). Variant callers to detect mutations include freebayes (red), MuSE (light green), Somatic Sniper (light blue), VarScan (pink), GATK (orange), Mutect2 (dark green), and Strelka2 (purple).*

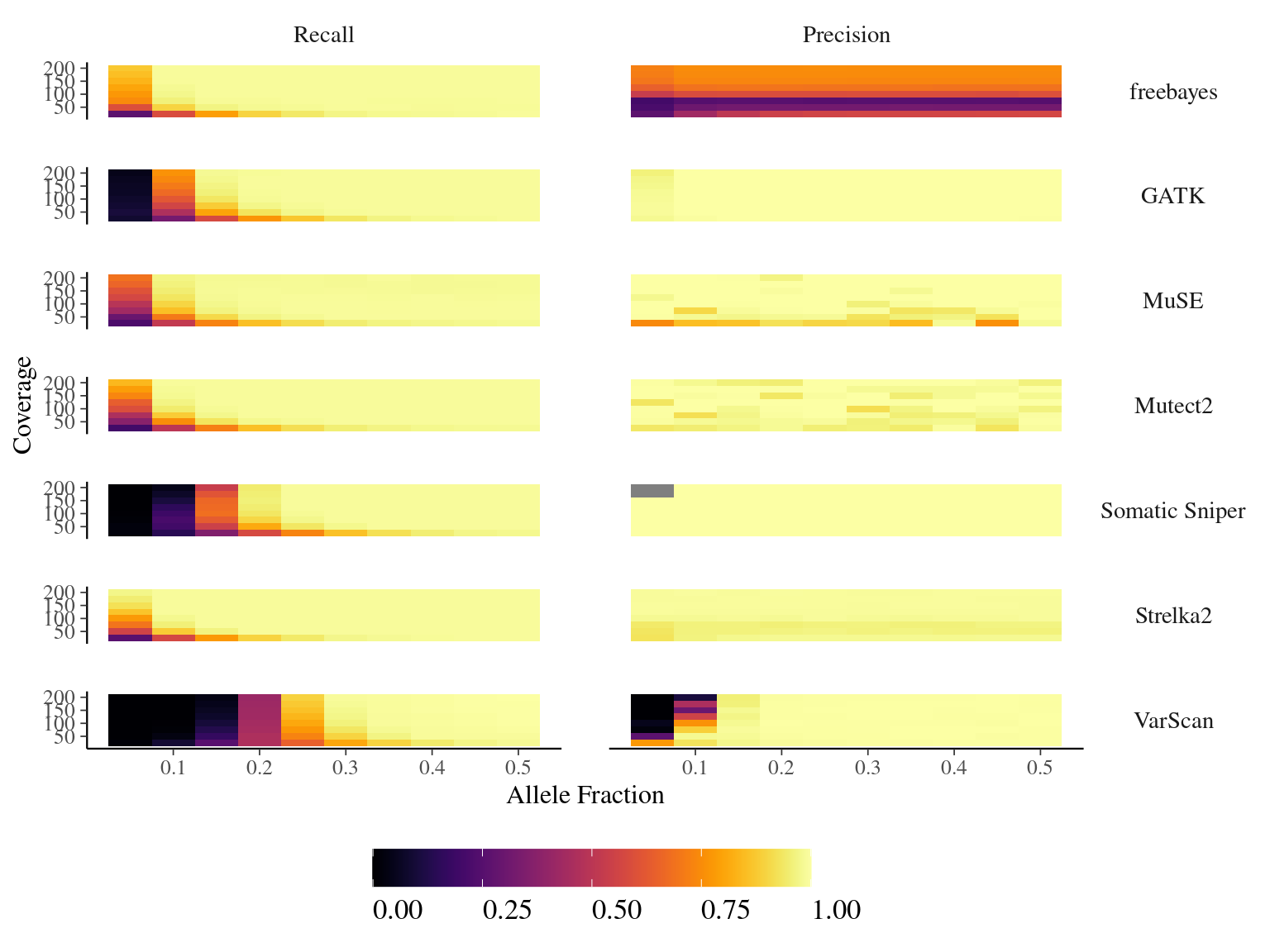

*Figure S3: Variation in the performance of variant callers for mutation detection with varying biological and sequencing parameters. The recall and the precision rates have been assessed for each variant caller to detect mutations with varying allelic fraction, and coverage (sequencing depth). Variant callers to detect mutations include freebayes (red), MuSE (light green), Somatic Sniper (light blue), VarScan (pink), GATK (orange), Mutect2 (dark green), and Strelka2 (purple).*

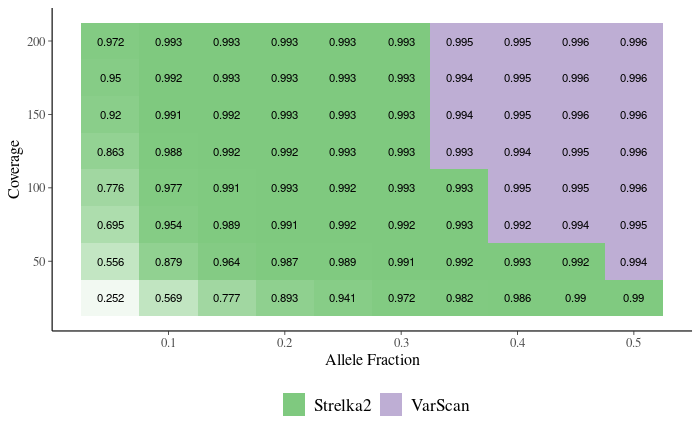

*Figure S4: Best performing variant callers for mutation detection depending on allelic fraction (allelic fraction) and coverage (sequencing depth). The best performing variant callers to detect mutations were Strelka2 (green) and VarScan (purple) for varying allelic fractions (x-axis) and sequencing depths (y-axis).*

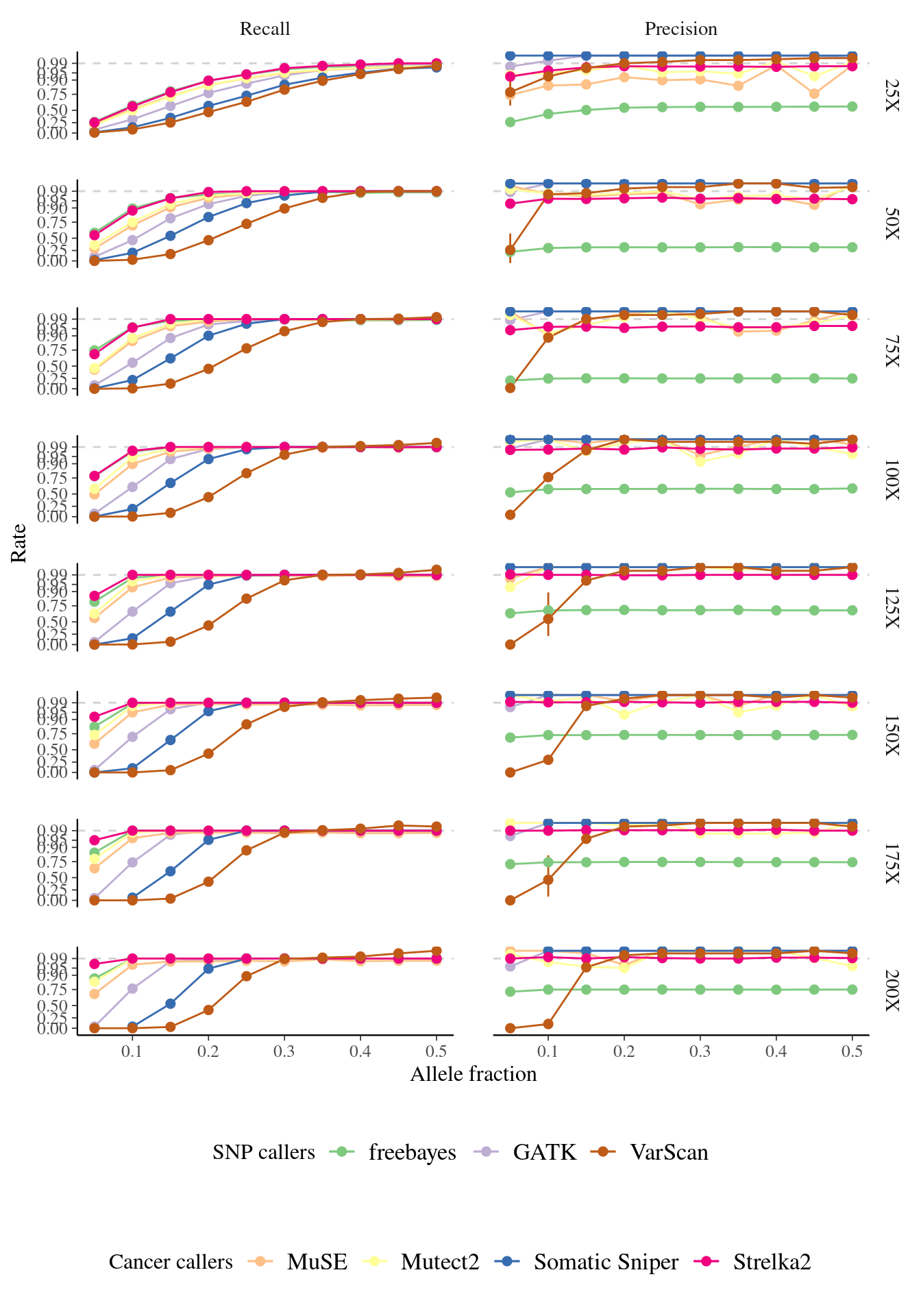

*Figure S5: Mutation recall and precision rates for SNP and cancer callers by allelic fraction and sequencing depth. Points represent mutation recall or precision (panel) rates (y-axis) for a given allelic fraction (x-axis) and sequencing depth (panel), with color representing variant callers: freebayes (green), GATK (light blue), VarScan (brown), MuSE (orange), Mutect2 (yellow), Somatic Sniper (dark blue), and Strelka2 (pink). The vertical lines represent the standard deviation of recall and precision rates over the 10 replicates. The light gray dashed line indicates a recall rate of 99%.*

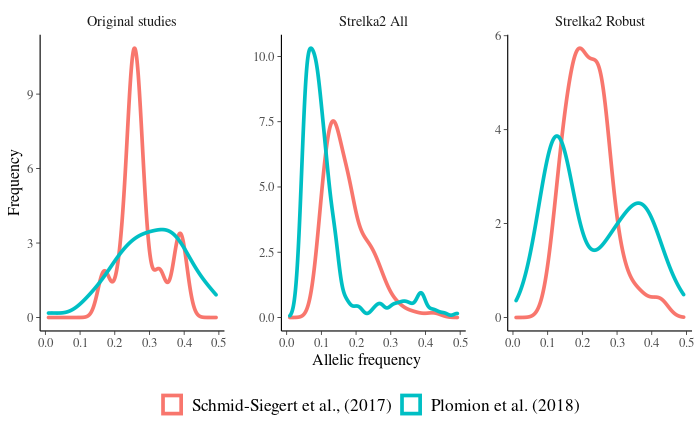

*Figure S6: Observed allelic frequencies of candidate mutations depending on variant callers and filtering in Schmid-Siegert et al., (2017) and Plomion et al., (2018). The datasets include the candidate mutations from the original article recovered in the raw data from the reanalysis, and Strelka2 after filtering, and the results of Strelka2 using the optimized filtering named robust.*

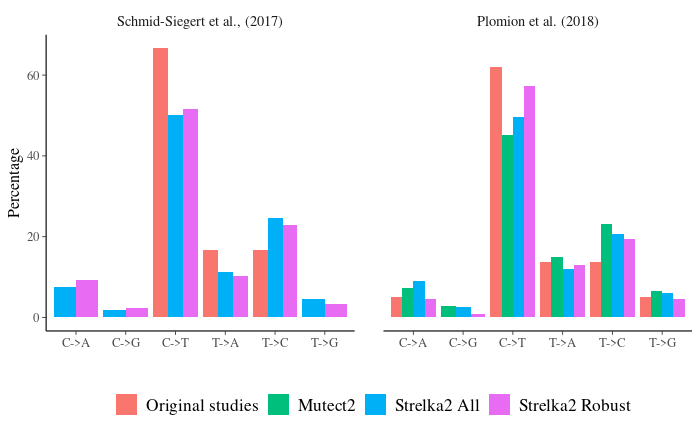

*Figure S7: Percentage of nucleotides change types of candidate mutations depending on variant callers and filtering in Schmid-Siegert et al., (2017) and Plomion et al., (2018). The datasets include the candidate mutations from the original article recovered in the raw data from the reanalysis (red), Mutect2 (green) and Strelka2 after filtering (blue), and the results of Strelka2 using the optimized filtering named robust (pink).*

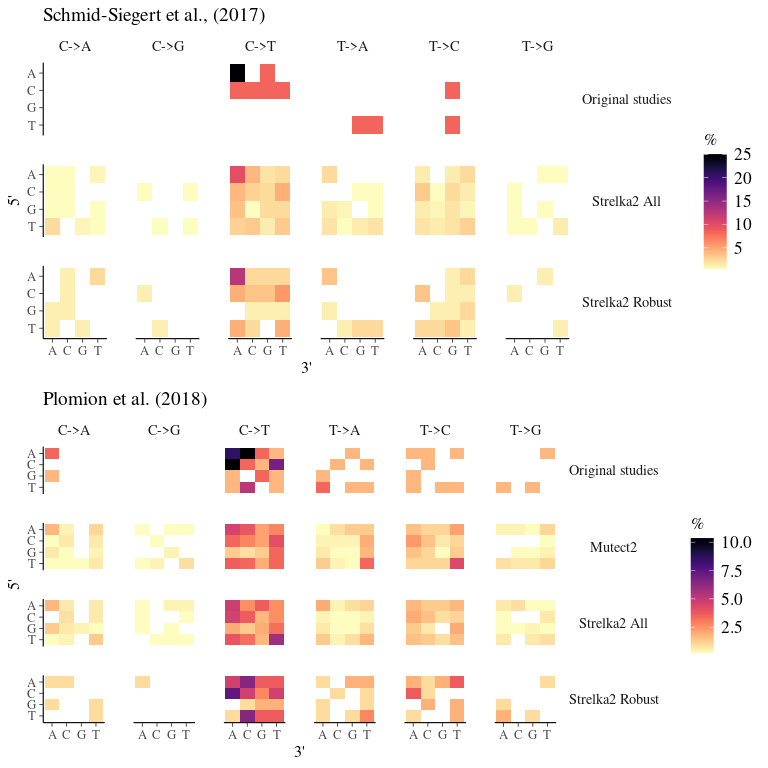

*Figure S8: Context-dependent mutation spectrum depending on variant callers and filtering in Schmid-Siegert et al., (2017) and Plomion et al., (2018). Heat maps show relative rates per triplet of nucleotides.*

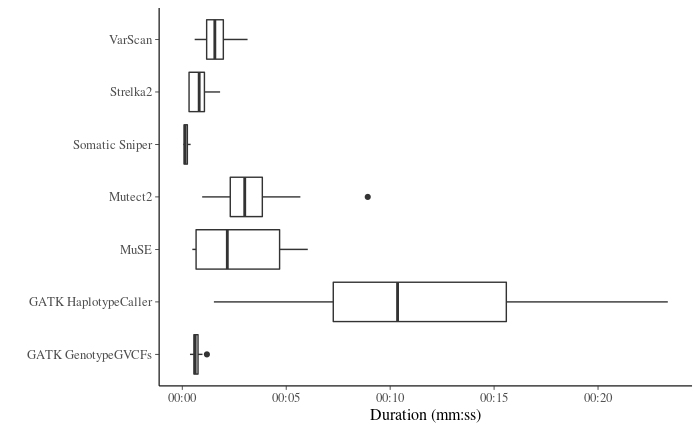

*Figure S9: Call duration over ten repetitions for each caller using 4 cpus and 16 GB of memory to detect a thousand mutations on a megabase. GATK duration is the addition of both HaplotypeCaller and GenotypeGVCFs. Mutect2, GATK, and VarScan did not include an option for parallelization and ran on 1 cpu.*

*Table S1: Correlations between simulations on the first megabase from the genome and ran five independent investigations based on randomly selected genome areas of a megabase in length.*

| **Caller** | **Recall** | **Precision** |
| --- | --- | --- |
| *freebayes* | 0.9991257 | 0.9976103 |
| *GATK* | 0.9995603 | 0.8449071 |
| *MuSE* | 0.9975487 | 0.3565296 |
| *Mutect2* | 0.9988121 | 0.7486251 |
| *Somatic Sniper* | 0.9993837 |  |
| *Strelka2* | 0.9981020 | 0.9450064 |
| *VarScan* | 0.9992663 | 0.8149302 |

*Table S2: Mean and standard deviation in performance of variant callers for mutation detection across all simulations. The recall and the precision rates have been assessed for each variant caller to detect mutations with varying allelic fraction, and sequencing depth.*

| **Caller** | **Recall** | **Precision** |
| --- | --- | --- |
| *freebayes* | 0.95 (0.11) | 0.57 (0.19) |
| *GATK* | 0.84 (0.29) | 1 (0.01) |
| *MuSE* | 0.91 (0.16) | 0.98 (0.11) |
| *Mutect2* | 0.93 (0.15) | 0.98 (0.07) |
| *Somatic Sniper* | 0.75 (0.36) | 1 (0) |
| *Strelka2* | 0.95 (0.11) | 0.98 (0.02) |
| *VarScan* | 0.63 (0.42) | 0.93 (0.23) |

*Table S3: Mean and standard deviation in performance of variant callers for mutation detection with varying allelic fraction and sequencing depth. The recall and the precision rates have been assessed for each variant caller to detect mutations with varying allelic fraction, and sequencing depth.*

| **Caller** | **Allelic fraction** | **Coverage** | **Recall** | **Precision** |
| --- | --- | --- | --- | --- |
| freebayes | 0.05 | 25 | 0.26 (0.01) | 0.26 (0.02) |
| freebayes | 0.05 | 50 | 0.59 (0.01) | 0.22 (0) |
| freebayes | 0.05 | 75 | 0.75 (0.01) | 0.2 (0.01) |
| freebayes | 0.05 | 100 | 0.77 (0.01) | 0.53 (0.01) |
| freebayes | 0.05 | 125 | 0.8 (0.02) | 0.64 (0.01) |
| freebayes | 0.05 | 150 | 0.83 (0.01) | 0.7 (0) |
| freebayes | 0.05 | 175 | 0.85 (0.01) | 0.72 (0.01) |
| freebayes | 0.05 | 200 | 0.87 (0.01) | 0.72 (0) |
| freebayes | 0.1 | 25 | 0.59 (0.02) | 0.43 (0.01) |
| freebayes | 0.1 | 50 | 0.9 (0.01) | 0.3 (0.01) |
| freebayes | 0.1 | 75 | 0.96 (0.01) | 0.25 (0.01) |
| freebayes | 0.1 | 100 | 0.97 (0.01) | 0.58 (0.01) |
| freebayes | 0.1 | 125 | 0.98 (0) | 0.69 (0.01) |
| freebayes | 0.1 | 150 | 0.99 (0) | 0.73 (0) |
| freebayes | 0.1 | 175 | 0.99 (0) | 0.74 (0.01) |
| freebayes | 0.1 | 200 | 0.99 (0) | 0.75 (0) |
| freebayes | 0.15 | 25 | 0.79 (0.01) | 0.51 (0.01) |
| freebayes | 0.15 | 50 | 0.96 (0.01) | 0.32 (0.01) |
| freebayes | 0.15 | 75 | 0.99 (0.01) | 0.25 (0) |
| freebayes | 0.15 | 100 | 0.99 (0) | 0.59 (0.01) |
| freebayes | 0.15 | 125 | 0.99 (0) | 0.69 (0.01) |
| freebayes | 0.15 | 150 | 0.99 (0) | 0.73 (0) |
| freebayes | 0.15 | 175 | 0.99 (0) | 0.74 (0) |
| freebayes | 0.15 | 200 | 0.99 (0) | 0.75 (0) |
| freebayes | 0.2 | 25 | 0.9 (0.01) | 0.55 (0.01) |
| freebayes | 0.2 | 50 | 0.98 (0) | 0.32 (0.01) |
| freebayes | 0.2 | 75 | 0.99 (0.01) | 0.25 (0) |
| freebayes | 0.2 | 100 | 0.99 (0) | 0.59 (0.01) |
| freebayes | 0.2 | 125 | 0.99 (0) | 0.69 (0.01) |
| freebayes | 0.2 | 150 | 0.99 (0) | 0.73 (0.01) |
| freebayes | 0.2 | 175 | 0.99 (0) | 0.75 (0.01) |
| freebayes | 0.2 | 200 | 0.99 (0) | 0.75 (0) |
| freebayes | 0.25 | 25 | 0.94 (0.01) | 0.56 (0.01) |
| freebayes | 0.25 | 50 | 0.98 (0.01) | 0.32 (0.01) |
| freebayes | 0.25 | 75 | 0.99 (0) | 0.25 (0) |
| freebayes | 0.25 | 100 | 0.99 (0) | 0.59 (0.01) |
| freebayes | 0.25 | 125 | 0.99 (0) | 0.69 (0.01) |
| freebayes | 0.25 | 150 | 0.99 (0) | 0.73 (0.01) |
| freebayes | 0.25 | 175 | 0.99 (0) | 0.75 (0.01) |
| freebayes | 0.25 | 200 | 0.99 (0) | 0.75 (0) |
| freebayes | 0.3 | 25 | 0.97 (0.01) | 0.56 (0.01) |
| freebayes | 0.3 | 50 | 0.99 (0) | 0.32 (0.01) |
| freebayes | 0.3 | 75 | 0.99 (0) | 0.25 (0.01) |
| freebayes | 0.3 | 100 | 0.99 (0) | 0.59 (0.01) |
| freebayes | 0.3 | 125 | 0.99 (0) | 0.69 (0.01) |
| freebayes | 0.3 | 150 | 0.99 (0) | 0.73 (0.01) |
| freebayes | 0.3 | 175 | 0.99 (0) | 0.75 (0.01) |
| freebayes | 0.3 | 200 | 0.99 (0) | 0.75 (0) |
| freebayes | 0.35 | 25 | 0.98 (0) | 0.56 (0.01) |
| freebayes | 0.35 | 50 | 0.99 (0) | 0.32 (0.01) |
| freebayes | 0.35 | 75 | 0.99 (0.01) | 0.25 (0) |
| freebayes | 0.35 | 100 | 0.99 (0) | 0.59 (0.01) |
| freebayes | 0.35 | 125 | 0.99 (0) | 0.69 (0.01) |
| freebayes | 0.35 | 150 | 0.99 (0) | 0.73 (0) |
| freebayes | 0.35 | 175 | 0.99 (0) | 0.74 (0.01) |
| freebayes | 0.35 | 200 | 0.99 (0) | 0.75 (0) |
| freebayes | 0.4 | 25 | 0.98 (0) | 0.56 (0.01) |
| freebayes | 0.4 | 50 | 0.99 (0.01) | 0.32 (0.01) |
| freebayes | 0.4 | 75 | 0.99 (0.01) | 0.25 (0.01) |
| freebayes | 0.4 | 100 | 0.99 (0) | 0.59 (0.01) |
| freebayes | 0.4 | 125 | 0.99 (0) | 0.69 (0) |
| freebayes | 0.4 | 150 | 0.99 (0) | 0.73 (0.01) |
| freebayes | 0.4 | 175 | 0.99 (0) | 0.74 (0.01) |
| freebayes | 0.4 | 200 | 0.99 (0) | 0.75 (0) |
| freebayes | 0.45 | 25 | 0.98 (0.01) | 0.57 (0.01) |
| freebayes | 0.45 | 50 | 0.99 (0.01) | 0.32 (0) |
| freebayes | 0.45 | 75 | 0.99 (0.01) | 0.25 (0) |
| freebayes | 0.45 | 100 | 0.99 (0) | 0.59 (0.01) |
| freebayes | 0.45 | 125 | 0.99 (0) | 0.69 (0.01) |
| freebayes | 0.45 | 150 | 0.99 (0) | 0.73 (0.01) |
| freebayes | 0.45 | 175 | 0.99 (0) | 0.74 (0.01) |
| freebayes | 0.45 | 200 | 0.99 (0) | 0.75 (0) |
| freebayes | 0.5 | 25 | 0.99 (0) | 0.57 (0.01) |
| freebayes | 0.5 | 50 | 0.99 (0.01) | 0.32 (0.01) |
| freebayes | 0.5 | 75 | 0.99 (0) | 0.25 (0.01) |
| freebayes | 0.5 | 100 | 0.99 (0) | 0.6 (0.02) |
| freebayes | 0.5 | 125 | 0.99 (0) | 0.69 (0.01) |
| freebayes | 0.5 | 150 | 0.99 (0) | 0.73 (0.01) |
| freebayes | 0.5 | 175 | 0.99 (0) | 0.74 (0) |
| freebayes | 0.5 | 200 | 0.99 (0) | 0.75 (0) |
| GATK | 0.05 | 25 | 0.08 (0.01) | 0.98 (0.02) |
| GATK | 0.05 | 50 | 0.1 (0.01) | 0.99 (0) |
| GATK | 0.05 | 75 | 0.09 (0.01) | 0.99 (0) |
| GATK | 0.05 | 100 | 0.08 (0.01) | 0.99 (0.01) |
| GATK | 0.05 | 125 | 0.06 (0.01) | 0.98 (0.01) |
| GATK | 0.05 | 150 | 0.06 (0.01) | 0.98 (0.01) |
| GATK | 0.05 | 175 | 0.06 (0.01) | 0.97 (0.01) |
| GATK | 0.05 | 200 | 0.04 (0.01) | 0.96 (0.01) |
| GATK | 0.1 | 25 | 0.32 (0.02) | 1 (0.01) |
| GATK | 0.1 | 50 | 0.46 (0.02) | 1 (0) |
| GATK | 0.1 | 75 | 0.56 (0.02) | 1 (0) |
| GATK | 0.1 | 100 | 0.62 (0.01) | 1 (0) |
| GATK | 0.1 | 125 | 0.67 (0.01) | 1 (0) |
| GATK | 0.1 | 150 | 0.71 (0.01) | 1 (0) |
| GATK | 0.1 | 175 | 0.74 (0.02) | 1 (0) |
| GATK | 0.1 | 200 | 0.76 (0.01) | 1 (0) |
| GATK | 0.15 | 25 | 0.58 (0.01) | 1 (0) |
| GATK | 0.15 | 50 | 0.8 (0.01) | 1 (0) |
| GATK | 0.15 | 75 | 0.88 (0.01) | 1 (0) |
| GATK | 0.15 | 100 | 0.93 (0.01) | 1 (0) |
| GATK | 0.15 | 125 | 0.96 (0.01) | 1 (0) |
| GATK | 0.15 | 150 | 0.97 (0.01) | 1 (0) |
| GATK | 0.15 | 175 | 0.98 (0.01) | 1 (0) |
| GATK | 0.15 | 200 | 0.98 (0) | 1 (0) |
| GATK | 0.2 | 25 | 0.77 (0.02) | 1 (0) |
| GATK | 0.2 | 50 | 0.93 (0.01) | 1 (0) |
| GATK | 0.2 | 75 | 0.97 (0.01) | 1 (0) |
| GATK | 0.2 | 100 | 0.98 (0.01) | 1 (0) |
| GATK | 0.2 | 125 | 0.99 (0.01) | 1 (0) |
| GATK | 0.2 | 150 | 0.99 (0) | 1 (0) |
| GATK | 0.2 | 175 | 0.99 (0) | 1 (0) |
| GATK | 0.2 | 200 | 0.99 (0.01) | 1 (0) |
| GATK | 0.25 | 25 | 0.87 (0.02) | 1 (0) |
| GATK | 0.25 | 50 | 0.97 (0.01) | 1 (0) |
| GATK | 0.25 | 75 | 0.98 (0.01) | 1 (0) |
| GATK | 0.25 | 100 | 0.99 (0) | 1 (0) |
| GATK | 0.25 | 125 | 0.99 (0) | 1 (0) |
| GATK | 0.25 | 150 | 0.99 (0) | 1 (0) |
| GATK | 0.25 | 175 | 0.99 (0) | 1 (0) |
| GATK | 0.25 | 200 | 0.99 (0) | 1 (0) |
| GATK | 0.3 | 25 | 0.93 (0.01) | 1 (0) |
| GATK | 0.3 | 50 | 0.99 (0.01) | 1 (0) |
| GATK | 0.3 | 75 | 0.99 (0) | 1 (0) |
| GATK | 0.3 | 100 | 0.99 (0) | 1 (0) |
| GATK | 0.3 | 125 | 0.99 (0) | 1 (0) |
| GATK | 0.3 | 150 | 0.99 (0) | 1 (0) |
| GATK | 0.3 | 175 | 0.99 (0) | 1 (0) |
| GATK | 0.3 | 200 | 0.99 (0) | 1 (0) |
| GATK | 0.35 | 25 | 0.97 (0.01) | 1 (0) |
| GATK | 0.35 | 50 | 0.99 (0) | 1 (0) |
| GATK | 0.35 | 75 | 0.99 (0) | 1 (0) |
| GATK | 0.35 | 100 | 0.99 (0) | 1 (0) |
| GATK | 0.35 | 125 | 0.99 (0) | 1 (0) |
| GATK | 0.35 | 150 | 0.99 (0) | 1 (0) |
| GATK | 0.35 | 175 | 0.99 (0) | 1 (0) |
| GATK | 0.35 | 200 | 0.99 (0) | 1 (0) |
| GATK | 0.4 | 25 | 0.98 (0) | 1 (0) |
| GATK | 0.4 | 50 | 0.99 (0) | 1 (0) |
| GATK | 0.4 | 75 | 0.99 (0) | 1 (0) |
| GATK | 0.4 | 100 | 0.99 (0) | 1 (0) |
| GATK | 0.4 | 125 | 0.99 (0) | 1 (0) |
| GATK | 0.4 | 150 | 0.99 (0) | 1 (0) |
| GATK | 0.4 | 175 | 0.99 (0) | 1 (0) |
| GATK | 0.4 | 200 | 0.99 (0) | 1 (0) |
| GATK | 0.45 | 25 | 0.98 (0.01) | 1 (0) |
| GATK | 0.45 | 50 | 0.99 (0) | 1 (0) |
| GATK | 0.45 | 75 | 0.99 (0) | 1 (0) |
| GATK | 0.45 | 100 | 0.99 (0) | 1 (0) |
| GATK | 0.45 | 125 | 0.99 (0) | 1 (0) |
| GATK | 0.45 | 150 | 0.99 (0) | 1 (0) |
| GATK | 0.45 | 175 | 0.99 (0) | 1 (0) |
| GATK | 0.45 | 200 | 0.99 (0) | 1 (0) |
| GATK | 0.5 | 25 | 0.99 (0) | 1 (0) |
| GATK | 0.5 | 50 | 0.99 (0) | 1 (0) |
| GATK | 0.5 | 75 | 0.99 (0) | 1 (0) |
| GATK | 0.5 | 100 | 0.99 (0) | 1 (0) |
| GATK | 0.5 | 125 | 0.99 (0) | 1 (0) |
| GATK | 0.5 | 150 | 0.99 (0) | 1 (0) |
| GATK | 0.5 | 175 | 0.99 (0) | 1 (0) |
| GATK | 0.5 | 200 | 0.99 (0) | 1 (0) |
| MuSE | 0.05 | 25 | 0.22 (0.01) | 0.74 (0.3) |
| MuSE | 0.05 | 50 | 0.3 (0.01) | 1 (0) |
| MuSE | 0.05 | 75 | 0.43 (0.02) | 1 (0) |
| MuSE | 0.05 | 100 | 0.49 (0.02) | 1 (0) |
| MuSE | 0.05 | 125 | 0.57 (0.01) | 0.98 (0.07) |
| MuSE | 0.05 | 150 | 0.61 (0.02) | 1 (0) |
| MuSE | 0.05 | 175 | 0.66 (0.02) | 1 (0) |
| MuSE | 0.05 | 200 | 0.69 (0.01) | 1 (0) |
| MuSE | 0.1 | 25 | 0.52 (0.02) | 0.85 (0.28) |
| MuSE | 0.1 | 50 | 0.71 (0.01) | 0.98 (0.04) |
| MuSE | 0.1 | 75 | 0.85 (0.02) | 0.9 (0.28) |
| MuSE | 0.1 | 100 | 0.9 (0.01) | 1 (0) |
| MuSE | 0.1 | 125 | 0.93 (0.01) | 1 (0) |
| MuSE | 0.1 | 150 | 0.95 (0.01) | 1 (0) |
| MuSE | 0.1 | 175 | 0.96 (0.01) | 1 (0) |
| MuSE | 0.1 | 200 | 0.97 (0) | 1 (0) |
| MuSE | 0.15 | 25 | 0.73 (0.01) | 0.86 (0.23) |
| MuSE | 0.15 | 50 | 0.91 (0.01) | 0.97 (0.06) |
| MuSE | 0.15 | 75 | 0.96 (0.01) | 0.99 (0.03) |
| MuSE | 0.15 | 100 | 0.97 (0.01) | 1 (0) |
| MuSE | 0.15 | 125 | 0.98 (0) | 1 (0) |
| MuSE | 0.15 | 150 | 0.98 (0.01) | 1 (0) |
| MuSE | 0.15 | 175 | 0.98 (0.01) | 1 (0) |
| MuSE | 0.15 | 200 | 0.98 (0) | 1 (0) |
| MuSE | 0.2 | 25 | 0.85 (0.02) | 0.92 (0.18) |
| MuSE | 0.2 | 50 | 0.97 (0.01) | 0.98 (0.05) |
| MuSE | 0.2 | 75 | 0.98 (0) | 1 (0) |
| MuSE | 0.2 | 100 | 0.99 (0.01) | 1 (0) |
| MuSE | 0.2 | 125 | 0.99 (0.01) | 1 (0) |
| MuSE | 0.2 | 150 | 0.99 (0.01) | 0.99 (0.02) |
| MuSE | 0.2 | 175 | 0.98 (0.01) | 1 (0) |
| MuSE | 0.2 | 200 | 0.98 (0) | 0.97 (0.1) |
| MuSE | 0.25 | 25 | 0.91 (0.01) | 0.9 (0.27) |
| MuSE | 0.25 | 50 | 0.98 (0) | 0.98 (0.04) |
| MuSE | 0.25 | 75 | 0.99 (0) | 1 (0) |
| MuSE | 0.25 | 100 | 0.99 (0) | 1 (0) |
| MuSE | 0.25 | 125 | 0.99 (0) | 1 (0) |
| MuSE | 0.25 | 150 | 0.98 (0.01) | 1 (0) |
| MuSE | 0.25 | 175 | 0.98 (0.01) | 1 (0) |
| MuSE | 0.25 | 200 | 0.98 (0.01) | 1 (0) |
| MuSE | 0.3 | 25 | 0.95 (0.01) | 0.91 (0.26) |
| MuSE | 0.3 | 50 | 0.99 (0) | 0.93 (0.21) |
| MuSE | 0.3 | 75 | 0.99 (0) | 1 (0.01) |
| MuSE | 0.3 | 100 | 0.99 (0) | 0.96 (0.11) |
| MuSE | 0.3 | 125 | 0.99 (0) | 1 (0) |
| MuSE | 0.3 | 150 | 0.99 (0.01) | 1 (0) |
| MuSE | 0.3 | 175 | 0.98 (0) | 1 (0) |
| MuSE | 0.3 | 200 | 0.98 (0) | 1 (0) |
| MuSE | 0.35 | 25 | 0.96 (0.01) | 0.85 (0.29) |
| MuSE | 0.35 | 50 | 0.99 (0) | 0.96 (0.1) |
| MuSE | 0.35 | 75 | 0.99 (0) | 0.93 (0.21) |
| MuSE | 0.35 | 100 | 0.99 (0) | 0.99 (0.03) |
| MuSE | 0.35 | 125 | 0.99 (0.01) | 1 (0) |
| MuSE | 0.35 | 150 | 0.98 (0.01) | 0.98 (0.06) |
| MuSE | 0.35 | 175 | 0.98 (0.01) | 1 (0) |
| MuSE | 0.35 | 200 | 0.99 (0) | 1 (0) |
| MuSE | 0.4 | 25 | 0.97 (0.01) | 0.98 (0.03) |
| MuSE | 0.4 | 50 | 0.99 (0) | 0.96 (0.11) |
| MuSE | 0.4 | 75 | 0.99 (0) | 0.94 (0.19) |
| MuSE | 0.4 | 100 | 0.99 (0) | 1 (0) |
| MuSE | 0.4 | 125 | 0.99 (0) | 1 (0) |
| MuSE | 0.4 | 150 | 0.98 (0) | 1 (0) |
| MuSE | 0.4 | 175 | 0.98 (0) | 1 (0) |
| MuSE | 0.4 | 200 | 0.98 (0) | 1 (0) |
| MuSE | 0.45 | 25 | 0.98 (0) | 0.76 (0.27) |
| MuSE | 0.45 | 50 | 0.99 (0) | 0.92 (0.23) |
| MuSE | 0.45 | 75 | 0.99 (0) | 0.99 (0.04) |
| MuSE | 0.45 | 100 | 0.99 (0) | 1 (0) |
| MuSE | 0.45 | 125 | 0.99 (0.01) | 1 (0) |
| MuSE | 0.45 | 150 | 0.98 (0.01) | 1 (0) |
| MuSE | 0.45 | 175 | 0.98 (0) | 1 (0) |
| MuSE | 0.45 | 200 | 0.98 (0.01) | 1 (0) |
| MuSE | 0.5 | 25 | 0.98 (0.01) | 0.98 (0.03) |
| MuSE | 0.5 | 50 | 0.99 (0) | 1 (0) |
| MuSE | 0.5 | 75 | 0.99 (0) | 1 (0) |
| MuSE | 0.5 | 100 | 0.99 (0) | 1 (0.01) |
| MuSE | 0.5 | 125 | 0.99 (0.01) | 1 (0) |
| MuSE | 0.5 | 150 | 0.98 (0.01) | 1 (0) |
| MuSE | 0.5 | 175 | 0.98 (0.01) | 1 (0) |
| MuSE | 0.5 | 200 | 0.98 (0.01) | 1 (0) |
| Mutect 2 | 0.05 | 25 | 0.2 (0.01) | 0.94 (0.06) |
| Mutect 2 | 0.05 | 50 | 0.37 (0.04) | 0.99 (0.01) |
| Mutect 2 | 0.05 | 75 | 0.46 (0.03) | 1 (0) |
| Mutect 2 | 0.05 | 100 | 0.59 (0.03) | 1 (0) |
| Mutect 2 | 0.05 | 125 | 0.64 (0.02) | 0.93 (0.18) |
| Mutect 2 | 0.05 | 150 | 0.73 (0.02) | 1 (0) |
| Mutect 2 | 0.05 | 175 | 0.78 (0.02) | 1 (0) |
| Mutect 2 | 0.05 | 200 | 0.84 (0.02) | 1 (0.01) |
| Mutect 2 | 0.1 | 25 | 0.5 (0.02) | 0.95 (0.09) |
| Mutect 2 | 0.1 | 50 | 0.75 (0.02) | 0.98 (0.03) |
| Mutect 2 | 0.1 | 75 | 0.88 (0.02) | 0.92 (0.17) |
| Mutect 2 | 0.1 | 100 | 0.95 (0.01) | 1 (0) |
| Mutect 2 | 0.1 | 125 | 0.97 (0.01) | 1 (0) |
| Mutect 2 | 0.1 | 150 | 0.98 (0) | 0.99 (0.01) |
| Mutect 2 | 0.1 | 175 | 0.98 (0.01) | 1 (0) |
| Mutect 2 | 0.1 | 200 | 0.99 (0) | 0.98 (0.06) |
| Mutect 2 | 0.15 | 25 | 0.72 (0.01) | 0.96 (0.05) |
| Mutect 2 | 0.15 | 50 | 0.93 (0.01) | 0.98 (0.05) |
| Mutect 2 | 0.15 | 75 | 0.97 (0.01) | 0.97 (0.06) |
| Mutect 2 | 0.15 | 100 | 0.98 (0.01) | 0.98 (0.05) |
| Mutect 2 | 0.15 | 125 | 0.99 (0) | 1 (0) |
| Mutect 2 | 0.15 | 150 | 0.99 (0) | 1 (0) |
| Mutect 2 | 0.15 | 175 | 0.99 (0) | 1 (0) |
| Mutect 2 | 0.15 | 200 | 0.99 (0) | 0.96 (0.13) |
| Mutect 2 | 0.2 | 25 | 0.86 (0.01) | 0.98 (0.04) |
| Mutect 2 | 0.2 | 50 | 0.98 (0.01) | 0.98 (0.04) |
| Mutect 2 | 0.2 | 75 | 0.99 (0) | 0.99 (0.02) |
| Mutect 2 | 0.2 | 100 | 0.99 (0) | 1 (0.01) |
| Mutect 2 | 0.2 | 125 | 0.99 (0) | 1 (0) |
| Mutect 2 | 0.2 | 150 | 0.99 (0) | 0.94 (0.14) |
| Mutect 2 | 0.2 | 175 | 0.99 (0) | 1 (0) |
| Mutect 2 | 0.2 | 200 | 0.99 (0) | 0.95 (0.15) |
| Mutect 2 | 0.25 | 25 | 0.92 (0.01) | 0.95 (0.12) |
| Mutect 2 | 0.25 | 50 | 0.98 (0.01) | 0.99 (0.03) |
| Mutect 2 | 0.25 | 75 | 0.99 (0) | 1 (0) |
| Mutect 2 | 0.25 | 100 | 0.99 (0) | 1 (0) |
| Mutect 2 | 0.25 | 125 | 0.99 (0) | 1 (0) |
| Mutect 2 | 0.25 | 150 | 0.99 (0) | 0.99 (0.02) |
| Mutect 2 | 0.25 | 175 | 0.99 (0) | 1 (0) |
| Mutect 2 | 0.25 | 200 | 0.99 (0) | 1 (0) |
| Mutect 2 | 0.3 | 25 | 0.95 (0.01) | 0.96 (0.1) |
| Mutect 2 | 0.3 | 50 | 0.99 (0) | 0.96 (0.12) |
| Mutect 2 | 0.3 | 75 | 0.99 (0) | 0.99 (0.03) |
| Mutect 2 | 0.3 | 100 | 0.99 (0) | 0.92 (0.15) |
| Mutect 2 | 0.3 | 125 | 0.99 (0) | 1 (0) |
| Mutect 2 | 0.3 | 150 | 0.99 (0) | 1 (0) |
| Mutect 2 | 0.3 | 175 | 0.99 (0) | 0.98 (0.04) |
| Mutect 2 | 0.3 | 200 | 0.99 (0) | 1 (0) |
| Mutect 2 | 0.35 | 25 | 0.97 (0.01) | 0.94 (0.11) |
| Mutect 2 | 0.35 | 50 | 0.99 (0) | 0.97 (0.06) |
| Mutect 2 | 0.35 | 75 | 0.99 (0) | 0.95 (0.14) |
| Mutect 2 | 0.35 | 100 | 0.99 (0) | 0.97 (0.1) |
| Mutect 2 | 0.35 | 125 | 0.99 (0) | 1 (0) |
| Mutect 2 | 0.35 | 150 | 0.99 (0) | 0.95 (0.15) |
| Mutect 2 | 0.35 | 175 | 0.99 (0) | 0.98 (0.05) |
| Mutect 2 | 0.35 | 200 | 0.99 (0) | 1 (0) |
| Mutect 2 | 0.4 | 25 | 0.98 (0.01) | 0.99 (0.01) |
| Mutect 2 | 0.4 | 50 | 0.99 (0) | 0.98 (0.07) |
| Mutect 2 | 0.4 | 75 | 0.99 (0) | 0.96 (0.13) |
| Mutect 2 | 0.4 | 100 | 0.99 (0) | 1 (0) |
| Mutect 2 | 0.4 | 125 | 0.99 (0) | 1 (0) |
| Mutect 2 | 0.4 | 150 | 0.99 (0) | 0.98 (0.06) |
| Mutect 2 | 0.4 | 175 | 0.99 (0) | 0.98 (0.06) |
| Mutect 2 | 0.4 | 200 | 0.99 (0) | 1 (0) |
| Mutect 2 | 0.45 | 25 | 0.98 (0) | 0.93 (0.1) |
| Mutect 2 | 0.45 | 50 | 0.99 (0) | 0.96 (0.13) |
| Mutect 2 | 0.45 | 75 | 0.99 (0) | 0.98 (0.07) |
| Mutect 2 | 0.45 | 100 | 0.99 (0) | 0.99 (0.03) |
| Mutect 2 | 0.45 | 125 | 0.99 (0) | 1 (0) |
| Mutect 2 | 0.45 | 150 | 0.99 (0) | 1 (0) |
| Mutect 2 | 0.45 | 175 | 0.99 (0) | 0.98 (0.05) |
| Mutect 2 | 0.45 | 200 | 0.99 (0) | 0.99 (0.02) |
| Mutect 2 | 0.5 | 25 | 0.98 (0) | 0.99 (0.01) |
| Mutect 2 | 0.5 | 50 | 0.99 (0) | 1 (0) |
| Mutect 2 | 0.5 | 75 | 0.99 (0) | 1 (0.01) |
| Mutect 2 | 0.5 | 100 | 0.99 (0) | 0.96 (0.08) |
| Mutect 2 | 0.5 | 125 | 0.99 (0) | 1 (0) |
| Mutect 2 | 0.5 | 150 | 0.99 (0) | 0.98 (0.06) |
| Mutect 2 | 0.5 | 175 | 0.99 (0) | 1 (0) |
| Mutect 2 | 0.5 | 200 | 0.99 (0) | 0.96 (0.12) |
| Somatic Sniper | 0.05 | 25 | 0.02 (0) | 1 (0) |
| Somatic Sniper | 0.05 | 50 | 0.02 (0.01) | 1 (0) |
| Somatic Sniper | 0.05 | 75 | 0.01 (0) | 1 (0) |
| Somatic Sniper | 0.05 | 100 | 0 (0) | 1 (0) |
| Somatic Sniper | 0.05 | 125 | 0 (0) | 1 (0) |
| Somatic Sniper | 0.05 | 150 | 0 (0) | 1 (0) |
| Somatic Sniper | 0.05 | 175 | 0 (0) | NaN (NA) |
| Somatic Sniper | 0.05 | 200 | 0 (0) | NaN (NA) |
| Somatic Sniper | 0.1 | 25 | 0.14 (0.01) | 1 (0) |
| Somatic Sniper | 0.1 | 50 | 0.2 (0.01) | 1 (0) |
| Somatic Sniper | 0.1 | 75 | 0.21 (0.01) | 1 (0) |
| Somatic Sniper | 0.1 | 100 | 0.19 (0.01) | 1 (0) |
| Somatic Sniper | 0.1 | 125 | 0.16 (0.01) | 1 (0) |
| Somatic Sniper | 0.1 | 150 | 0.11 (0.01) | 1 (0) |
| Somatic Sniper | 0.1 | 175 | 0.07 (0.01) | 1 (0) |
| Somatic Sniper | 0.1 | 200 | 0.04 (0) | 1 (0) |
| Somatic Sniper | 0.15 | 25 | 0.35 (0.01) | 1 (0) |
| Somatic Sniper | 0.15 | 50 | 0.54 (0.02) | 1 (0) |
| Somatic Sniper | 0.15 | 75 | 0.63 (0.02) | 1 (0) |
| Somatic Sniper | 0.15 | 100 | 0.68 (0.01) | 1 (0) |
| Somatic Sniper | 0.15 | 125 | 0.67 (0.01) | 1 (0) |
| Somatic Sniper | 0.15 | 150 | 0.66 (0.01) | 1 (0) |
| Somatic Sniper | 0.15 | 175 | 0.61 (0.02) | 1 (0) |
| Somatic Sniper | 0.15 | 200 | 0.53 (0.01) | 1 (0) |
| Somatic Sniper | 0.2 | 25 | 0.58 (0.02) | 1 (0) |
| Somatic Sniper | 0.2 | 50 | 0.81 (0.02) | 1 (0) |
| Somatic Sniper | 0.2 | 75 | 0.9 (0.01) | 1 (0) |
| Somatic Sniper | 0.2 | 100 | 0.93 (0.01) | 1 (0) |
| Somatic Sniper | 0.2 | 125 | 0.95 (0.01) | 1 (0) |
| Somatic Sniper | 0.2 | 150 | 0.96 (0.01) | 1 (0) |
| Somatic Sniper | 0.2 | 175 | 0.95 (0.01) | 1 (0) |
| Somatic Sniper | 0.2 | 200 | 0.95 (0.01) | 1 (0) |
| Somatic Sniper | 0.25 | 25 | 0.73 (0.01) | 1 (0) |
| Somatic Sniper | 0.25 | 50 | 0.94 (0.01) | 1 (0) |
| Somatic Sniper | 0.25 | 75 | 0.97 (0.01) | 1 (0) |
| Somatic Sniper | 0.25 | 100 | 0.98 (0.01) | 1 (0) |
| Somatic Sniper | 0.25 | 125 | 0.99 (0) | 1 (0) |
| Somatic Sniper | 0.25 | 150 | 0.99 (0) | 1 (0) |
| Somatic Sniper | 0.25 | 175 | 0.99 (0) | 1 (0) |
| Somatic Sniper | 0.25 | 200 | 0.99 (0) | 1 (0) |
| Somatic Sniper | 0.3 | 25 | 0.86 (0.01) | 1 (0) |
| Somatic Sniper | 0.3 | 50 | 0.98 (0.01) | 1 (0) |
| Somatic Sniper | 0.3 | 75 | 0.99 (0) | 1 (0) |
| Somatic Sniper | 0.3 | 100 | 0.99 (0) | 1 (0) |
| Somatic Sniper | 0.3 | 125 | 0.99 (0) | 1 (0) |
| Somatic Sniper | 0.3 | 150 | 0.99 (0) | 1 (0) |
| Somatic Sniper | 0.3 | 175 | 0.99 (0) | 1 (0) |
| Somatic Sniper | 0.3 | 200 | 0.99 (0) | 1 (0) |
| Somatic Sniper | 0.35 | 25 | 0.92 (0.01) | 1 (0) |
| Somatic Sniper | 0.35 | 50 | 0.99 (0) | 1 (0) |
| Somatic Sniper | 0.35 | 75 | 0.99 (0) | 1 (0) |
| Somatic Sniper | 0.35 | 100 | 0.99 (0) | 1 (0) |
| Somatic Sniper | 0.35 | 125 | 0.99 (0) | 1 (0) |
| Somatic Sniper | 0.35 | 150 | 0.99 (0) | 1 (0) |
| Somatic Sniper | 0.35 | 175 | 0.99 (0) | 1 (0) |
| Somatic Sniper | 0.35 | 200 | 0.99 (0) | 1 (0) |
| Somatic Sniper | 0.4 | 25 | 0.95 (0.01) | 1 (0) |
| Somatic Sniper | 0.4 | 50 | 0.99 (0) | 1 (0) |
| Somatic Sniper | 0.4 | 75 | 0.99 (0) | 1 (0) |
| Somatic Sniper | 0.4 | 100 | 0.99 (0) | 1 (0) |
| Somatic Sniper | 0.4 | 125 | 0.99 (0) | 1 (0) |
| Somatic Sniper | 0.4 | 150 | 0.99 (0) | 1 (0) |
| Somatic Sniper | 0.4 | 175 | 0.99 (0) | 1 (0) |
| Somatic Sniper | 0.4 | 200 | 0.99 (0) | 1 (0) |
| Somatic Sniper | 0.45 | 25 | 0.97 (0.01) | 1 (0) |
| Somatic Sniper | 0.45 | 50 | 0.99 (0) | 1 (0) |
| Somatic Sniper | 0.45 | 75 | 0.99 (0) | 1 (0) |
| Somatic Sniper | 0.45 | 100 | 0.99 (0) | 1 (0) |
| Somatic Sniper | 0.45 | 125 | 0.99 (0) | 1 (0) |
| Somatic Sniper | 0.45 | 150 | 0.99 (0) | 1 (0) |
| Somatic Sniper | 0.45 | 175 | 0.99 (0) | 1 (0) |
| Somatic Sniper | 0.45 | 200 | 0.99 (0) | 1 (0) |
| Somatic Sniper | 0.5 | 25 | 0.98 (0.01) | 1 (0) |
| Somatic Sniper | 0.5 | 50 | 0.99 (0) | 1 (0) |
| Somatic Sniper | 0.5 | 75 | 0.99 (0) | 1 (0) |
| Somatic Sniper | 0.5 | 100 | 0.99 (0) | 1 (0) |
| Somatic Sniper | 0.5 | 125 | 0.99 (0) | 1 (0) |
| Somatic Sniper | 0.5 | 150 | 0.99 (0) | 1 (0) |
| Somatic Sniper | 0.5 | 175 | 0.99 (0) | 1 (0) |
| Somatic Sniper | 0.5 | 200 | 0.99 (0) | 1 (0) |
| Strelka 2 | 0.05 | 25 | 0.25 (0.01) | 0.93 (0.01) |
| Strelka 2 | 0.05 | 50 | 0.55 (0.01) | 0.93 (0.01) |
| Strelka 2 | 0.05 | 75 | 0.69 (0.02) | 0.94 (0.01) |
| Strelka 2 | 0.05 | 100 | 0.78 (0.01) | 0.98 (0.01) |
| Strelka 2 | 0.05 | 125 | 0.86 (0.01) | 0.99 (0.01) |
| Strelka 2 | 0.05 | 150 | 0.92 (0.01) | 0.99 (0) |
| Strelka 2 | 0.05 | 175 | 0.95 (0.01) | 0.99 (0) |
| Strelka 2 | 0.05 | 200 | 0.97 (0) | 0.99 (0) |
| Strelka 2 | 0.1 | 25 | 0.57 (0.02) | 0.96 (0.01) |
| Strelka 2 | 0.1 | 50 | 0.88 (0.01) | 0.96 (0.01) |
| Strelka 2 | 0.1 | 75 | 0.96 (0.01) | 0.96 (0) |
| Strelka 2 | 0.1 | 100 | 0.98 (0) | 0.98 (0) |
| Strelka 2 | 0.1 | 125 | 0.99 (0) | 0.99 (0) |
| Strelka 2 | 0.1 | 150 | 0.99 (0) | 0.99 (0) |
| Strelka 2 | 0.1 | 175 | 0.99 (0) | 0.99 (0) |
| Strelka 2 | 0.1 | 200 | 0.99 (0) | 0.99 (0.01) |
| Strelka 2 | 0.15 | 25 | 0.78 (0.01) | 0.98 (0.01) |
| Strelka 2 | 0.15 | 50 | 0.96 (0.01) | 0.96 (0.01) |
| Strelka 2 | 0.15 | 75 | 0.99 (0) | 0.96 (0.01) |
| Strelka 2 | 0.15 | 100 | 0.99 (0) | 0.99 (0.01) |
| Strelka 2 | 0.15 | 125 | 0.99 (0) | 0.99 (0) |
| Strelka 2 | 0.15 | 150 | 0.99 (0) | 0.99 (0) |
| Strelka 2 | 0.15 | 175 | 0.99 (0) | 0.99 (0) |
| Strelka 2 | 0.15 | 200 | 0.99 (0) | 0.99 (0) |
| Strelka 2 | 0.2 | 25 | 0.89 (0.01) | 0.98 (0.01) |
| Strelka 2 | 0.2 | 50 | 0.99 (0) | 0.96 (0.01) |
| Strelka 2 | 0.2 | 75 | 0.99 (0) | 0.95 (0.01) |
| Strelka 2 | 0.2 | 100 | 0.99 (0) | 0.98 (0) |
| Strelka 2 | 0.2 | 125 | 0.99 (0) | 0.99 (0) |
| Strelka 2 | 0.2 | 150 | 0.99 (0) | 0.99 (0) |
| Strelka 2 | 0.2 | 175 | 0.99 (0) | 0.99 (0) |
| Strelka 2 | 0.2 | 200 | 0.99 (0) | 0.99 (0.01) |
| Strelka 2 | 0.25 | 25 | 0.94 (0.01) | 0.98 (0) |
| Strelka 2 | 0.25 | 50 | 0.99 (0) | 0.97 (0.01) |
| Strelka 2 | 0.25 | 75 | 0.99 (0) | 0.96 (0.01) |
| Strelka 2 | 0.25 | 100 | 0.99 (0) | 0.99 (0) |
| Strelka 2 | 0.25 | 125 | 0.99 (0) | 0.99 (0) |
| Strelka 2 | 0.25 | 150 | 0.99 (0) | 0.99 (0) |
| Strelka 2 | 0.25 | 175 | 0.99 (0) | 0.99 (0) |
| Strelka 2 | 0.25 | 200 | 0.99 (0) | 0.99 (0) |
| Strelka 2 | 0.3 | 25 | 0.97 (0.01) | 0.98 (0) |
| Strelka 2 | 0.3 | 50 | 0.99 (0) | 0.96 (0.01) |
| Strelka 2 | 0.3 | 75 | 0.99 (0) | 0.96 (0) |
| Strelka 2 | 0.3 | 100 | 0.99 (0) | 0.98 (0.01) |
| Strelka 2 | 0.3 | 125 | 0.99 (0) | 0.99 (0) |
| Strelka 2 | 0.3 | 150 | 0.99 (0) | 0.99 (0) |
| Strelka 2 | 0.3 | 175 | 0.99 (0) | 0.99 (0) |
| Strelka 2 | 0.3 | 200 | 0.99 (0) | 0.99 (0) |
| Strelka 2 | 0.35 | 25 | 0.98 (0) | 0.98 (0.01) |
| Strelka 2 | 0.35 | 50 | 0.99 (0) | 0.96 (0.01) |
| Strelka 2 | 0.35 | 75 | 0.99 (0) | 0.96 (0.01) |
| Strelka 2 | 0.35 | 100 | 0.99 (0) | 0.98 (0.01) |
| Strelka 2 | 0.35 | 125 | 0.99 (0) | 0.99 (0.01) |
| Strelka 2 | 0.35 | 150 | 0.99 (0) | 0.99 (0) |
| Strelka 2 | 0.35 | 175 | 0.99 (0) | 0.99 (0) |
| Strelka 2 | 0.35 | 200 | 0.99 (0) | 0.99 (0) |
| Strelka 2 | 0.4 | 25 | 0.99 (0.01) | 0.98 (0) |
| Strelka 2 | 0.4 | 50 | 0.99 (0) | 0.96 (0.01) |
| Strelka 2 | 0.4 | 75 | 0.99 (0) | 0.96 (0) |
| Strelka 2 | 0.4 | 100 | 0.99 (0) | 0.99 (0.01) |
| Strelka 2 | 0.4 | 125 | 0.99 (0) | 0.99 (0.01) |
| Strelka 2 | 0.4 | 150 | 0.99 (0) | 0.99 (0) |
| Strelka 2 | 0.4 | 175 | 0.99 (0) | 0.99 (0) |
| Strelka 2 | 0.4 | 200 | 0.99 (0) | 0.99 (0) |
| Strelka 2 | 0.45 | 25 | 0.99 (0) | 0.98 (0) |
| Strelka 2 | 0.45 | 50 | 0.99 (0) | 0.96 (0.01) |
| Strelka 2 | 0.45 | 75 | 0.99 (0) | 0.96 (0.01) |
| Strelka 2 | 0.45 | 100 | 0.99 (0) | 0.99 (0.01) |
| Strelka 2 | 0.45 | 125 | 0.99 (0) | 0.99 (0) |
| Strelka 2 | 0.45 | 150 | 0.99 (0) | 0.99 (0) |
| Strelka 2 | 0.45 | 175 | 0.99 (0) | 0.99 (0) |
| Strelka 2 | 0.45 | 200 | 0.99 (0) | 0.99 (0) |
| Strelka 2 | 0.5 | 25 | 0.99 (0) | 0.98 (0) |
| Strelka 2 | 0.5 | 50 | 0.99 (0) | 0.96 (0.01) |
| Strelka 2 | 0.5 | 75 | 0.99 (0) | 0.96 (0.01) |
| Strelka 2 | 0.5 | 100 | 0.99 (0) | 0.99 (0) |
| Strelka 2 | 0.5 | 125 | 0.99 (0) | 0.99 (0) |
| Strelka 2 | 0.5 | 150 | 0.99 (0) | 0.99 (0.01) |
| Strelka 2 | 0.5 | 175 | 0.99 (0) | 0.99 (0) |
| Strelka 2 | 0.5 | 200 | 0.99 (0) | 0.99 (0) |
| VarScan | 0.05 | 25 | 0.01 (0) | 0.78 (0.19) |
| VarScan | 0.05 | 50 | 0 (0) | 0.26 (0.32) |
| VarScan | 0.05 | 75 | 0 (0) | 0.02 (0.05) |
| VarScan | 0.05 | 100 | 0 (0) | 0.05 (0.12) |
| VarScan | 0.05 | 125 | 0 (0) | 0 (NA) |
| VarScan | 0.05 | 150 | 0 (0) | 0 (0) |
| VarScan | 0.05 | 175 | 0 (0) | 0 (0) |
| VarScan | 0.05 | 200 | 0 (0) | 0 (0) |
| VarScan | 0.1 | 25 | 0.09 (0.02) | 0.93 (0.03) |
| VarScan | 0.1 | 50 | 0.03 (0) | 0.98 (0.04) |
| VarScan | 0.1 | 75 | 0.01 (0) | 0.89 (0.13) |
| VarScan | 0.1 | 100 | 0 (0.01) | 0.76 (0.32) |
| VarScan | 0.1 | 125 | 0 (0) | 0.55 (0.34) |
| VarScan | 0.1 | 150 | 0 (0) | 0.3 (0.04) |
| VarScan | 0.1 | 175 | 0 (0) | 0.46 (0.36) |
| VarScan | 0.1 | 200 | 0 (0) | 0.11 (0.13) |
| VarScan | 0.15 | 25 | 0.25 (0.01) | 0.97 (0.02) |
| VarScan | 0.15 | 50 | 0.17 (0.01) | 0.98 (0.01) |
| VarScan | 0.15 | 75 | 0.13 (0.01) | 0.99 (0.02) |
| VarScan | 0.15 | 100 | 0.1 (0.01) | 0.98 (0.02) |
| VarScan | 0.15 | 125 | 0.07 (0.01) | 0.97 (0.04) |
| VarScan | 0.15 | 150 | 0.06 (0.01) | 0.98 (0.02) |
| VarScan | 0.15 | 175 | 0.04 (0.01) | 0.96 (0.07) |
| VarScan | 0.15 | 200 | 0.03 (0.01) | 0.95 (0.05) |
| VarScan | 0.2 | 25 | 0.47 (0.02) | 0.99 (0.01) |
| VarScan | 0.2 | 50 | 0.46 (0.01) | 1 (0.01) |
| VarScan | 0.2 | 75 | 0.45 (0.02) | 1 (0) |
| VarScan | 0.2 | 100 | 0.44 (0.01) | 1 (0) |
| VarScan | 0.2 | 125 | 0.43 (0.01) | 1 (0) |
| VarScan | 0.2 | 150 | 0.43 (0.02) | 1 (0) |
| VarScan | 0.2 | 175 | 0.42 (0.02) | 1 (0.01) |
| VarScan | 0.2 | 200 | 0.42 (0.02) | 1 (0.01) |
| VarScan | 0.25 | 25 | 0.64 (0.02) | 0.99 (0.01) |
| VarScan | 0.25 | 50 | 0.73 (0.01) | 1 (0) |
| VarScan | 0.25 | 75 | 0.77 (0.01) | 1 (0) |
| VarScan | 0.25 | 100 | 0.81 (0.01) | 1 (0) |
| VarScan | 0.25 | 125 | 0.83 (0.01) | 1 (0) |
| VarScan | 0.25 | 150 | 0.86 (0.01) | 1 (0) |
| VarScan | 0.25 | 175 | 0.87 (0.01) | 1 (0) |
| VarScan | 0.25 | 200 | 0.89 (0.01) | 1 (0) |
| VarScan | 0.3 | 25 | 0.81 (0.01) | 1 (0.01) |
| VarScan | 0.3 | 50 | 0.9 (0.01) | 1 (0) |
| VarScan | 0.3 | 75 | 0.93 (0.01) | 1 (0) |
| VarScan | 0.3 | 100 | 0.96 (0.01) | 1 (0) |
| VarScan | 0.3 | 125 | 0.97 (0) | 1 (0) |
| VarScan | 0.3 | 150 | 0.98 (0.01) | 1 (0) |
| VarScan | 0.3 | 175 | 0.98 (0.01) | 1 (0) |
| VarScan | 0.3 | 200 | 0.99 (0) | 1 (0) |
| VarScan | 0.35 | 25 | 0.89 (0.01) | 1 (0.01) |
| VarScan | 0.35 | 50 | 0.97 (0.01) | 1 (0) |
| VarScan | 0.35 | 75 | 0.98 (0) | 1 (0) |
| VarScan | 0.35 | 100 | 0.99 (0) | 1 (0) |
| VarScan | 0.35 | 125 | 0.99 (0) | 1 (0) |
| VarScan | 0.35 | 150 | 0.99 (0) | 1 (0) |
| VarScan | 0.35 | 175 | 0.99 (0) | 1 (0) |
| VarScan | 0.35 | 200 | 0.99 (0) | 1 (0) |
| VarScan | 0.4 | 25 | 0.94 (0.01) | 1 (0) |
| VarScan | 0.4 | 50 | 0.99 (0) | 1 (0) |
| VarScan | 0.4 | 75 | 0.99 (0) | 1 (0) |
| VarScan | 0.4 | 100 | 0.99 (0) | 1 (0) |
| VarScan | 0.4 | 125 | 0.99 (0) | 1 (0) |
| VarScan | 0.4 | 150 | 1 (0.01) | 1 (0) |
| VarScan | 0.4 | 175 | 0.99 (0.01) | 1 (0) |
| VarScan | 0.4 | 200 | 0.99 (0.01) | 1 (0) |
| VarScan | 0.45 | 25 | 0.97 (0.01) | 1 (0) |
| VarScan | 0.45 | 50 | 0.99 (0) | 1 (0.01) |
| VarScan | 0.45 | 75 | 0.99 (0) | 1 (0) |
| VarScan | 0.45 | 100 | 0.99 (0.01) | 1 (0.01) |
| VarScan | 0.45 | 125 | 0.99 (0.01) | 1 (0) |
| VarScan | 0.45 | 150 | 1 (0) | 1 (0) |
| VarScan | 0.45 | 175 | 1 (0) | 1 (0) |
| VarScan | 0.45 | 200 | 1 (0) | 1 (0) |
| VarScan | 0.5 | 25 | 0.98 (0) | 1 (0) |
| VarScan | 0.5 | 50 | 0.99 (0) | 1 (0) |
| VarScan | 0.5 | 75 | 0.99 (0.01) | 1 (0) |
| VarScan | 0.5 | 100 | 1 (0) | 1 (0) |
| VarScan | 0.5 | 125 | 1 (0) | 1 (0) |
| VarScan | 0.5 | 150 | 1 (0) | 1 (0) |
| VarScan | 0.5 | 175 | 1 (0) | 1 (0) |
| VarScan | 0.5 | 200 | 1 (0) | 1 (0) |
